# Bacterial internalins exploit E-cadherin to promote head and neck tumor metastasis and drug resistance

**DOI:** 10.64898/2026.04.20.719623

**Authors:** Hongxia Wang, Christopher Pham, Kurni Kurniyati, Zheng Liu, Jinyang Cai, Michael J Lynch, Jiong Li, Claire D. James, Iain M. Morgan, Brian R Crane, Xiang-yang Wang, Chunhao Li

## Abstract

Head and neck squamous cell carcinoma (HNSCC) is an aggressive malignancy characterized by local invasion, lymph node metastasis, and therapeutic resistance. Chronic periodontal disease has been linked to HNSCC progression, yet the responsible pathogens and underlying molecular mechanisms remain unclear. Here, we show that the keystone periodontal pathogen *Porphyromonas gingivalis* promotes HNSCC metastasis and chemoresistance through two internalin proteins that are secreted via the type IX secretion system (T9SS). These internalin proteins specifically bind the EC1 domain of E-cadherin through their curved solenoid-like leucine-rich repeats (LRRs), facilitating bacterial invasion and inducing epithelial-to-mesenchymal transition (EMT). Mechanistically, internalin-E-cadherin engagement drives β-catenin nuclear translocation and activates p38 and JNK1/2 MAP kinase signaling pathways, enhancing tumor cell migration, metastatic dissemination, and resistance to cisplatin-induced apoptosis. Tissue microarrays detect internalin antigens in HNSCC specimens, supporting their in vivo relevance. Together, these findings establish a direct mechanistic link between an oral pathogen and HNSCC progression and extend the paradigm of internalin-E-cadherin interactions from microbial pathogenesis to cancer biology.

## INTRODUCTION

Oral squamous cell carcinoma (OSCC), the most common malignancy of the oral cavity and a major subset of head and neck squamous cell carcinoma (HNSCC), remains a significant global health challenge with poor overall survival rates^1,2^. High mortality is driven by aggressive local invasion, lymph node metastasis, and frequent therapeutic failure^3,4^. While tobacco use, alcohol consumption, and human papillomavirus (HPV) infection are well-established drivers of tumor initiation^2,3^, metastatic progression, the primary cause of mortality remains poorly understood and inadequately targeted therapeutically. Emerging evidence suggests that the oral microbiota is not merely a passive bystander in HNSCC but an active constituent of the tumor microenvironment that can shape malignant behavior^5–7^.

The oral cavity harbors one of the most densely populated and diverse microbiota in the human body, existing in constant contact with stratified squamous epithelia^8–10^. Microbial dysbiosis, particularly in the setting of chronic periodontitis, disrupts epithelial barrier integrity, sustains chronic inflammation, and increases cancer risk^9,11^. Recent spatial transcriptomic and single-cell sequencing analyses have revealed heterogeneous intratumoral microbial communities within HNSCC lesions, implicating direct host-microbe interactions within tumor tissues^5,12^. However, whether and how specific oral bacteria directly engage epithelial tumor cells to regulate invasive and metastatic behavior remains largely unexplored.

*Porphyromonas gingivalis*, one of the keystone pathogens of periodontitis, exerts disproportionate pathogenic effects on host immunity, tissue integrity, and microbial homeostasis^13,14^. Beyond its established role in periodontal disease, *P. gingivalis* is frequently enriched in OSCC tissues and has been associated with enhanced tumor cell survival and immune evasion^6,7,15^. Experimental infection models further demonstrate that *P. gingivalis* promotes tumor cell migration and invasion, supporting a causal role in metastatic progression^15–17^. Despite these observations, the bacterial effectors and host receptors that mediate these pro-tumorigenic activities remain undefined, limiting mechanistic insights and hindering therapeutic translation.

Disruption of epithelial cell-cell adhesion is a hallmark of tumor invasion and metastasis^18,19^. Adherens junctions (AJs), centered on the transmembrane adhesion molecule E-cadherin, are critical for maintaining epithelial architecture and tissue homeostasis. Loss or functional impairment of E-cadherin is strongly linked to invasive growth, epithelial-to-mesenchymal transition (EMT), and poor clinical outcomes in HNSCC and other solid tumors^20–23^. In addition to genetic and epigenetic alterations, E-cadherin function can be perturbed through direct interactions with microbial factors^24,25^. A canonical example is the internalin proteins of *Listeria monocytogenes*, a bacterial pathogen that causes listeriosis ^26^. This group of proteins typically harbor multiple leucine-rich repeats (LRRs) and bind E-cadherin to promote bacterial invasion into host cells^27–29^. Importantly, internalin-E-cadherin engagement not only facilitates bacterial entry but also destabilizes AJs and activates signaling pathways governing cytoskeletal remodeling and cell motility^30–32^. Despite these well-characterized roles in bacterial infections, the relevance of internalin-mediated E-cadherin modulation to tumorigenesis remains largely unexplored.

*P. gingivalis* utilizes a Type IX secretion system (T9SS) to export a diverse repertoire of surface-associated and secreted proteins, including gingipains, a family of cysteine proteinases that act as a primary virulence determinant in causing chronic periodontitis and systemic inflammation^33–36^. However, the biological functions of many T9SS cargo substrates remain poorly defined. Here, we combined bioinformatic analysis with functional screening to systematically interrogate T9SS-secreted proteins and identified two internalin-like proteins containing multiple LRRs. We demonstrate that these two internalins specifically bind human E-cadherin and facilitate bacterial invasion. Using complementary *in vitro* and *in vivo* models, we further show that internalin-E-cadherin interactions enhance HNSCC cell migration, invasion, metastatic dissemination, and resistance to anticancer therapy. Finally, we delineate downstream signaling pathways triggered by the engagement of E-cadherin-internalins. Collectively, these findings reveal a previously unrecognized mechanism by which an oral pathogen directly hijacks epithelial adhesion machinery to promote tumor metastasis, extending the relevance of internalin-E-cadherin interactions beyond classical infectious disease paradigms to cancer progression.

## RESULTS

### Identification of two internalin-like proteins toxic to yeast and mammalian cells

Approximately 30 proteins have been identified as T9SS cargo substrates in *P. gingivalis*^35,37^, yet most remain functionally uncharacterized. To systematically assess their biological activity, we screened these proteins for their toxicity in yeast and identified multiple candidates that impaired yeast growth to varying extents. Among these, PG0350 (484 aa, 52.7 kDa) was previously identified as an internalin-related protein that mediates interdomain community development between *P. gingivalis* and *Candida albicans* ^38^ and PG1374 (428 aa, 47.1 kDa) as an immunoreactive antigen^39^. These two proteins share ∼29% sequence identity and exhibit similar domain architectures, including a short N-terminal Sec-dependent signal peptide (SP), a β-strand-rich region, 7-9 leucine-rich repeats (LRRs), a linker, and a conserved C-terminal domain (CTD) that serves as a T9SS secretion signal^40^(**Figure 1A**). AlphaFold structural modeling indicated that the region containing LRRs adopts a curved solenoid-like conformation (**Figure 1B**), a characteristic of bacterial internalin proteins^27,41^. Yeast toxicity assays showed that expression of PG0350 or PG1374 markedly reduced yeast colony formation (**Figure 1C**). Consistently, transient expression of GFP-tagged PG0350 or PG1374 in HEK293T cells, but not GFP alone, induced cell rounding, shrinkage, and lysis (**Figure 1D-E**), indicating cytotoxicity in mammalian cells. Together, these data identify PG0350 and PG1374 as putative T9SS-secreted virulence factors with internalin-like structural features. Based on their shared LRR architectures to *L. monocytogenes* internalins^27^, we designate PG0350 and PG1374 as InlA and InlB, respectively.

**Figure 1.**
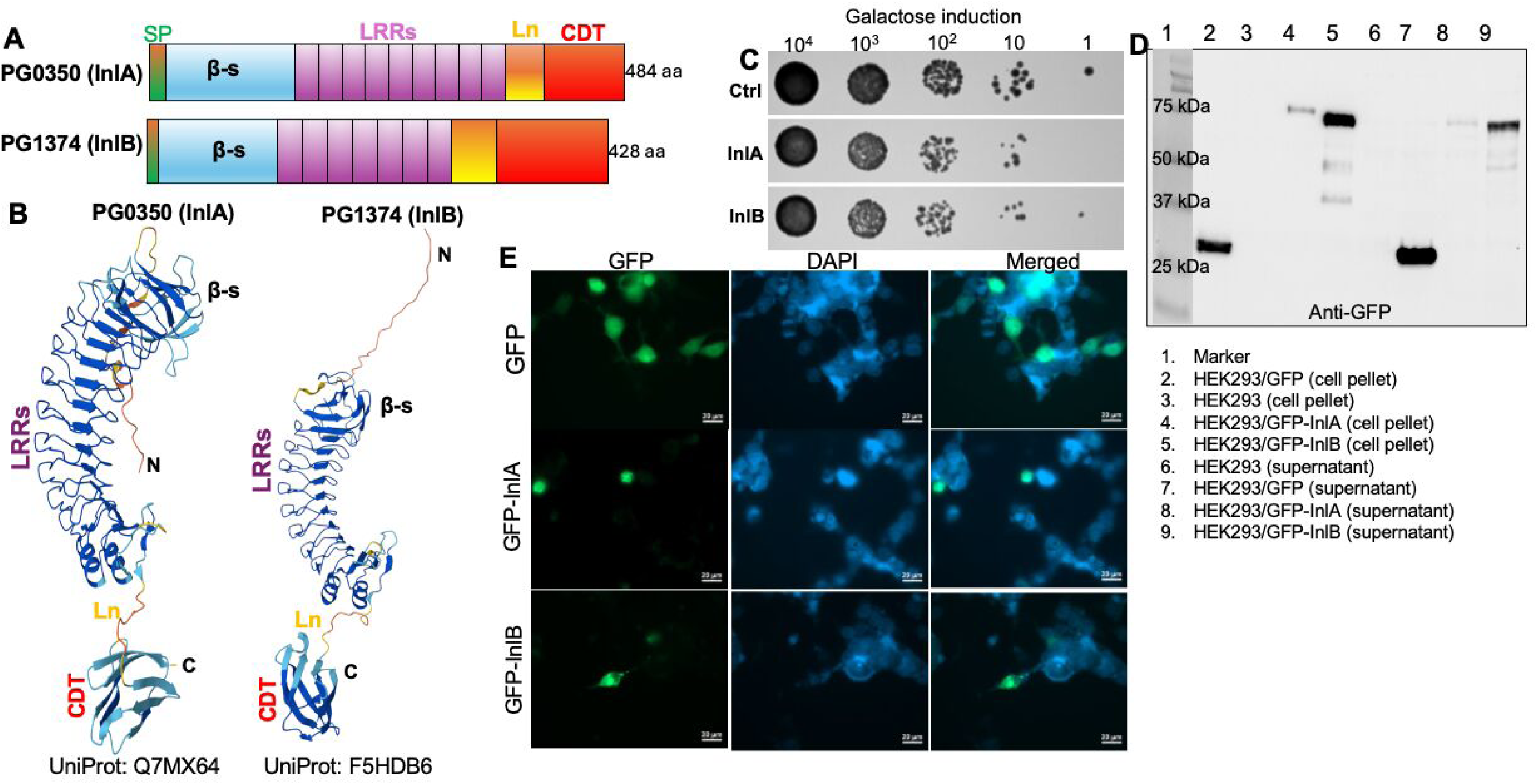
Identification of two internalin-like proteins in *P. gingivalis*. **(A)** Schematic diagrams illustrating the domain architecture of PG0350 (InlA) and PG1374 (InlB). SP, Sec-dependent signal peptide; β-s: β-strands; LRRs, leucine-rich repeats; Ln, linker; CTD, C-terminal domain. **(B)** AlphaFold structural models of InlA (UniProt: Q7MX64) and InlB (UniProt: F5HDB6). N, N-terminus; C, C-terminus. **(C)** InlA and InlB are toxic to yeast cells. Yeast strains expressing InlA (residues 26-484) or InlB (residues 24-428) under a galactose-inducible promoter (PGAL1; pYES2/NTA2 plasmid) were serially diluted and spotted onto plates containing glucose or galactose. Plates were incubated at 30 °C for 48 h prior to imaging. Ctrl, empty vector control. **(D-E)** InlA and InlB are toxic to mammalian cells. Codon-optimized InlA and InlB sequences were cloned into the pcDNA6.2 C-EmGFP-GW TOPO expression vector (Invitrogen) and transiently transfected into HEK293T cells. The expression of GFP-tagged InlA/InlB was confirmed by immunoblotting using anti-GFP antibody (D) and by fluorescence microscopy (E). Images were acquired 72 h post-transfection. Scale bars, 20 μm.

### InlA and InlB are T9SS-secreted and promote bacterial cell invasion

To investigate the roles of InlA and InlB, we generated deletion mutants (*ΔinlA* and *ΔinlB*) and their corresponding isogenic complemented strains (**Figure S1** & **S2**). These genetically modified strains were confirmed by PCR and immunoblotting using InlA- and InlB-specific antibodies (αInlA and αInlB). Deletion of either gene has no impact on *P. gingivalis* growth in TSB medium (**Figure S3**). To determine whether InlA and InlB are secreted via T9SS, we generated a *porT* deletion mutant (*ΔporT*), as PorT is essential for T9SS function^42^. Consistent with a T9SS-deficient phenotype, the *ΔporT* mutant formed beige instead of black-pigmented colonies on blood agar plates (**Figure S4**). Immunoblotting analysis detected InlA and InlB in both whole-cell lysates and culture supernatants of the wild-type W83 strain. In contrast, these proteins were detected only in the whole-cell lysates, but not in the supernatants of *ΔporT* mutant (**Figure 2A-B**). As expected, InlA and InlB were absent in their respective deletion mutants and restored in the complemented strains (**Figure 2A-B**). Notably, two immunoreactive bands were frequently observed in the whole-cell lysates of W83 and two complemented strains (**Figure 2; Figures S1** & **S2**). The sizes of these bands correspond to InlA and InlB proteins with or without CTD, consistent with differential processing during T9SS-mediated secretion.

**Figure 2.**
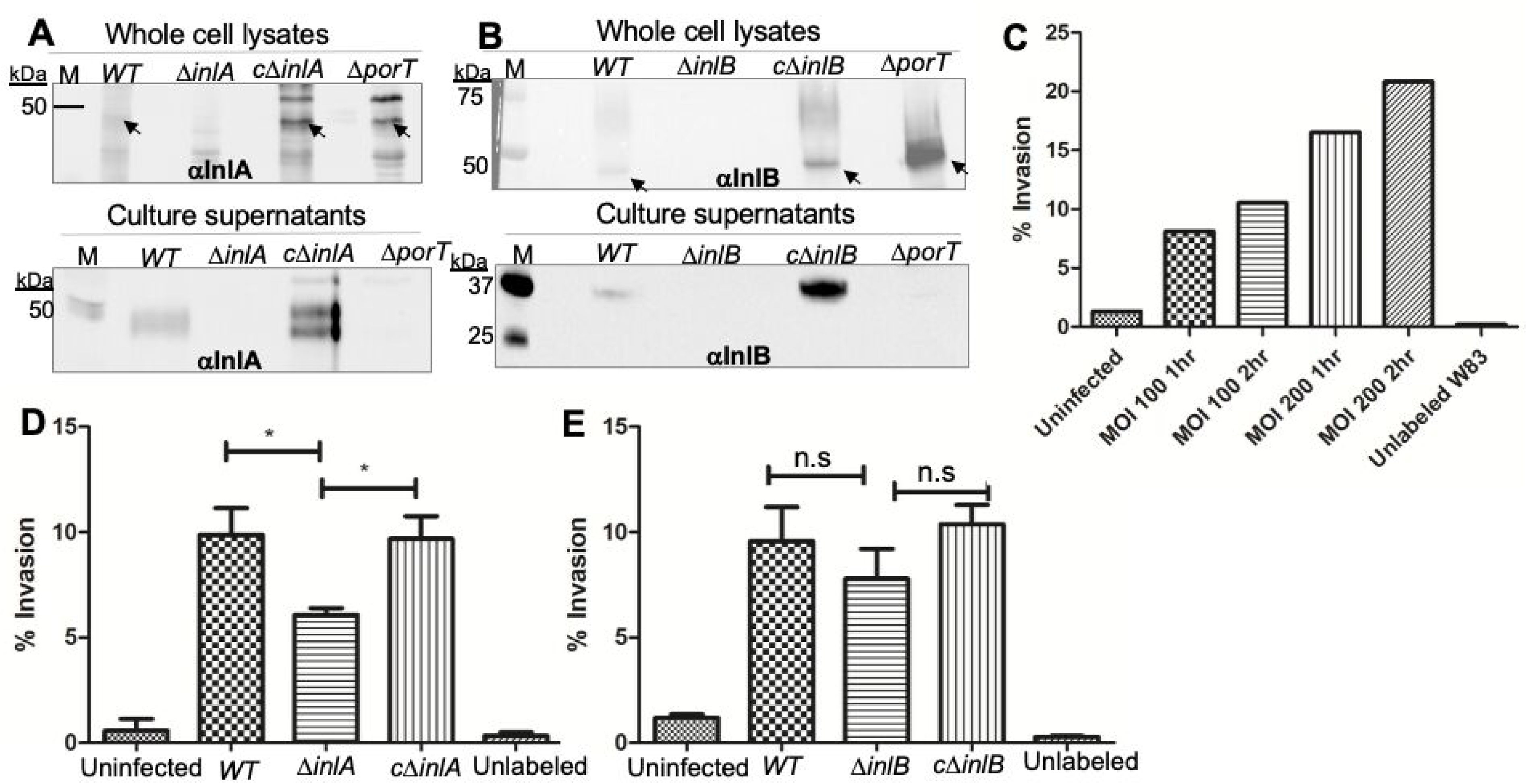
InlA and InlB are secreted via the T9SS and contribute to *P. gingivalis* invasion. **(A)** Immunoblot detection of InlA in whole-cell lysates (top) and culture supernatants (bottom) from WT W83, Δ*inlA*, complemented (cΔ*inlA*), and Δ*porT* strains using anti-InlA antibody (αInlA). **(B)** Immunoblot detection of InlB in whole-cell lysates (top) and culture supernatants (bottom) from WT, Δ*inlB*, complemented (cΔ*inlB*), and Δ*porT* strains using anti-InlB antibody (αInlB). **(C)** *P. gingivalis* invades TIGK cells in a dose-dependent manner. TIGK cells were infected with BCECF-AM-labeled or unlabeled *P. gingivalis* W83 at MOI 100, 200 for 1 and 2 h at 37°C under anaerobic conditions. After trypan blue quenching of extracellular fluorescence, bacterial internalization was quantified by flow cytometry. Percent invasion represents the proportion of cells containing internalized bacteria among 10,000 events (mean ± SEM; three independent experiments performed in duplicate). **(D-E)** InlA and InlB contribute to *P. gingivalis* invasion of host cells. TIGK cells were infected with BCECF-AM-labeled *P. gingivalis* W83, *ΔinlA* **(D)** or *ΔinlB* **(E)** mutants, and their respective complemented strains (*cΔinlA, cΔinlB*) at MOI 100 for 1 h at 37°C under anaerobic conditions. Percent invasion was measured as described in **(C)**. Data are presented as mean ± SEM. Statistical analysis was performed using one-way ANOVA with Tukey’s post hoc test; *\*P < 0.05*; N.S., not significant.

We next assessed whether InlA and InlB contribute to *P. gingivalis* cell invasion using human telomerase-immortalized gingival keratinocytes (TIGK). For this study, TIGK cells were infected with different bacterial strains as indicated, and intracellular bacteria were quantified by flow cytometry (**Figure S5**). We found that deletion of *inl*A significantly reduced bacterial invasion (**Figure 2D**), whereas *ΔinlB* showed a modest but not statistically significant decrease compared with W83 and the complemented strain (**Figure 2E**). To further define the kinetics of invasion, TIGK cells were infected with W83 at MOIs of 100 or 200 for 1 or 2 hours. Invasion increased in a dose-dependent manner at both time points, with approximately two-fold higher invasion at an MOI of 200 compared with 100 (8.1% vs 16.5% at 1 hour; 10.5% vs 20.8% at 2 hours). In contrast, extending infection time from 1 to 2 hours resulted in only modest increases at the same MOI (**Figure 2C**), indicating that invasion occurs largely within the first hour of infection. We also attempted to generate a double mutant lacking both *inlA* and *inlB* to evaluate potential functional overlap in cell invasion; however, these attempts were unsuccessful. Together, these data demonstrate that InlA and InlB are secreted through the T9SS and contribute to *P. gingivalis* cell invasion.

### InlA and InlB bind TIGK cells through E-cadherin

Because InlA and InlB promote *P. gingivalis* invasion of TIGK cells, we next sought to identify the host receptor mediating their binding. Co-immuno-precipitation (co-IP) assays using recombinant His-tagged InlA (rInlA) or InlB (rInlB) and TIGK cell lysates revealed that E-cadherin was specifically pulled down by both proteins using Ni-NTA beads, indicating a direct interaction (**Figure 3A-B**). This interaction was further confirmed by co-IP and ELISA using Fc-tagged E-cadherin (rE-cad) and rInlB (**Figure 3C; Figure S6A-C**). Consistent with the co-IP and ELISA data, immunofluorescence microscopy analysis (IFA) demonstrated colocalization of FITC-labeled InlA or InlB with E-cadherin on the TIGK cell surface (**Figure 3D**).

**Figure 3.**
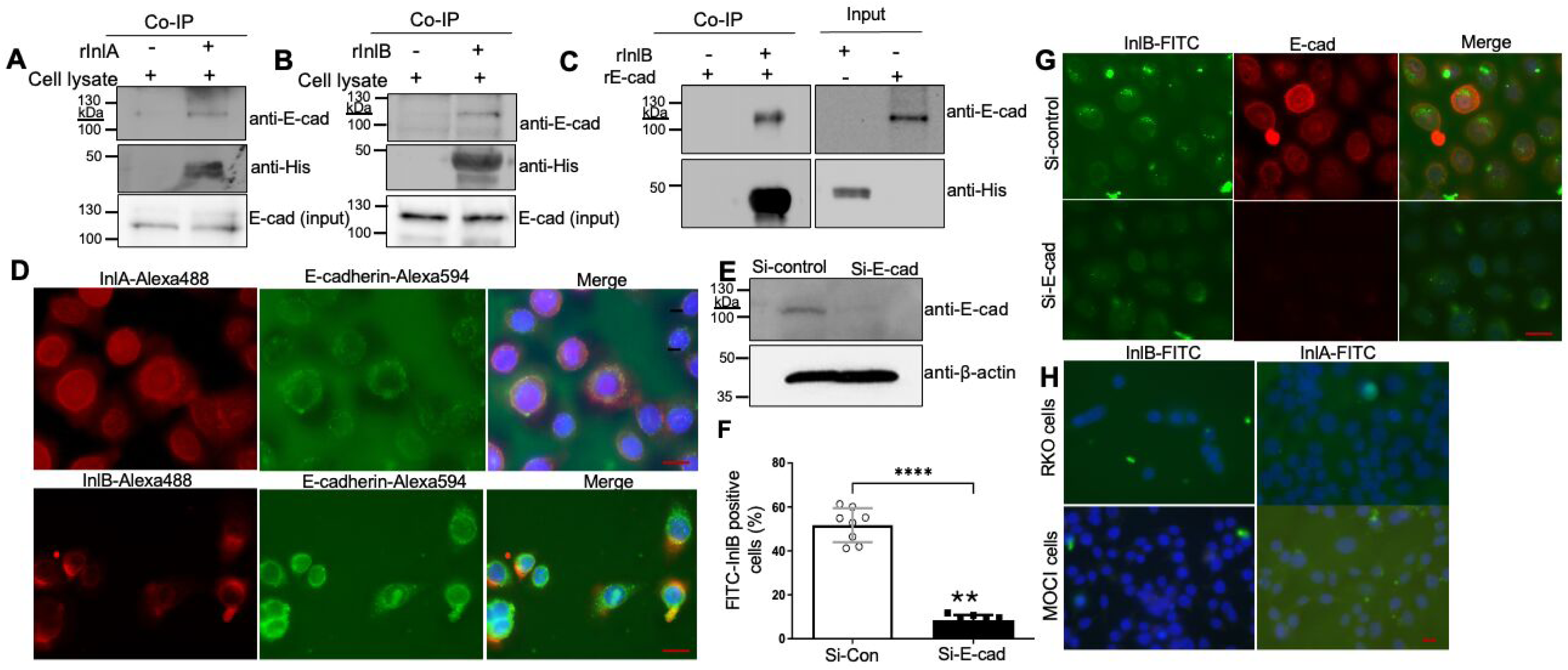
InlA and InlB bind TIGK cells through E-cadherin. **(A-B)** Co-immunoprecipitation (co-IP) assays. TIGK cell lysates were incubated with His-tagged recombinant InlA (rInlA) or InlB (rInlB; 20 μg/ml) for 3 hours at 4 °C, followed by incubation with Ni-NTA resin overnight. After washing (PBS, 0.05% Tween 20), bound proteins were eluted in Laemmli buffer and analyzed by immunoblotting with anti-His and anti-E-cadherin antibodies. **(C)** Co-IP assays using rInlB and recombinant Fc-E-cadherin (rE-cad). **(D)** Co-localization of InlA/InlB with E-cadherin. TIGK cells were incubated with FITC-labeled InlA or InlB (20 μg/ml, 2 h), followed by immunostaining with anti-E-cadherin and Alexa Fluor 594-conjugated secondary antibodies. Nuclei were counterstained with DAPI. Images were acquired using a Zeiss fluorescence microscope with a 63× oil-immersion objective. Scale bars: 20 μm. **(E)** E-cadherin knockdown efficiency in siRNA-transfected TIGK cells was confirmed by immunoblotting. **(F)** Quantification of FITC-InlB-positive cells was performed using ImageJ. **(G)** Binding of FITC-InlB to TIGK cells following E-cadherin knockdown was assessed by immuno-fluorescence microscopy. Scale bars: 20 μm. **(H)** Binding and internalization of FITC-labeled InlA or InlB in RKO and MOC1 cells were analyzed by immunofluorescence staining. Scale bars: 20 μm.

To assess the functional role of E-cadherin in protein binding, E-cadherin expression was silenced in TIGK cells using siRNA, with knockdown efficiency confirmed by immunoblots using a specific antibody against human E-cadherin (**Figure 3E**). E-cadherin depletion nearly abolished InlA and InlB binding to TIGK cells, as shown by IFA (**Figure 3G; Figure S6D**). Specifically, quantification analysis revealed a reduction in FITC-InlB-positive cells from ∼56% in control cells to <10% following E-cadherin knockdown (**Figure 3F**). Similar patterns were observed in HSC3^43^, a human tongue squamous carcinoma cell line (**Figure S6E**). Furthermore, FITC-labeled InlA or InlB did not bind RKO cells, which lack E-cadherin expression^44^, or MOC1 cells expressing murine E-cadherin^45^, demonstrating their binding specificity for human E-cadherin (**Figure 3H**). Collectively, these results establish human E-cadherin as a host receptor for InlA and InlB and mediates *P. gingivalis* binding and entry into gingival epithelial cells.

### InlA and InlB bind the Ec1 domain of E-cadherin with high affinity

The ectodomain (Ec) of E-cadherin comprises five tandem repeats (Ec1-Ec5; **Figure 4A**). To validate the interaction between InlA/InlB and E-cadherin, and to define their binding sites, we performed surface plasmon resonance (SPR) analysis using His-tagged InlA or InlB as immobilized ligands and GST-tagged E-cadherin constructs (Ec1-5, Ec1, Ec2-5, or the point mutant Ec1^P16E^) as analytes. All recombinant proteins were expressed in *E. coli* and purified using FPLC (**Figure 4B**). Ligand proteins were immobilized on SPR sensor chips, and binding kinetics were measured using increasing concentrations (0.5-10 µM) of GST-tagged analytes. InlB bound robustly to full-length E-cadherin ectodomain (Ec1-5; **Figure 4C-D**) and to Ec1 (**Figure 4E-F**), exhibiting high association rate constants (InlB/Ec1-5, *k*_a_ = 7.25 x 10^4^ M^-1^s^-1^; InlB/Ec1, *k*_a_ = 1.12 x 10^5^ M^-1^s^-1^) (see measurements in **Table S1**), indicative of high-affinity interactions. In contrast, no detectable binding was observed between InlB and Ec2-5, demonstrating that Ec1 is required for the interaction (**Figure 4G-H**).

**Figure 4.**
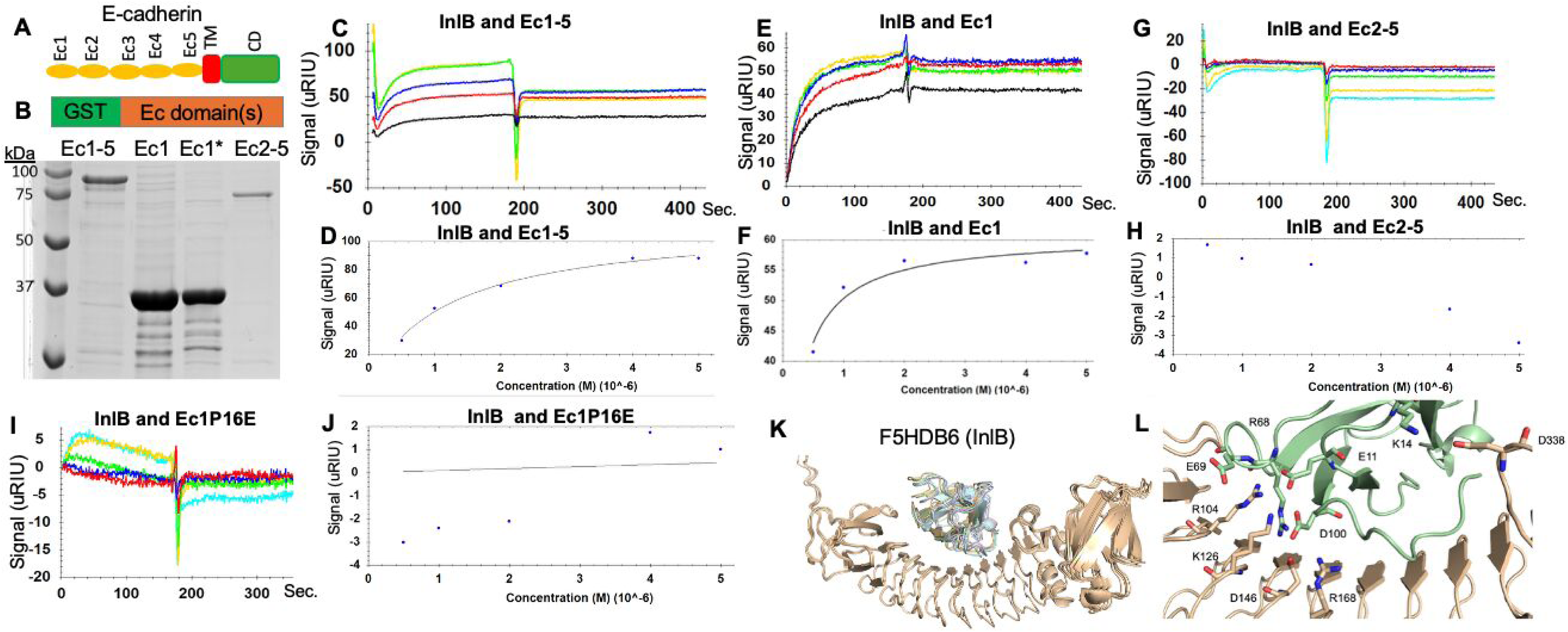
InlB binds the extracellular domains of E-cadherin with high affinity. **(A)** Schematic of E-cadherin showing five extracellular cadherin repeats (Ec1-5), transmembrane domain (TM), and cytoplasmic domain (CD). **(B)** SDS-PAGE of purified GST-tagged E-cadherin constructs: Ec1-5, Ec1, Ec1* (Ec1^P16E^), and Ec2-5. **(C, E, G, I)** SPR sensorgrams of InlB binding to Ec1-5 (C), Ec1 (E), Ec2-5 (G), and Ec1P16E (I) at multiple concentrations, showing association and dissociation kinetics. **(D, F, H, J)** Dose-dependent saturation binding of InlB to Ec1-5 (D), Ec1 (F), Ec2-5 (H), and Ec1^P16E^ (J). The equilibrium dissociation constant (K_D_) was calculated for each binding experiment. **(K)** Structural modeling shows that Ec1 (cyan) interacts in the internal grove of InlB (tan, F5HDB6). **(L)** Close-up of the InlB-Ec1 interface highlights key residues and hydrogen bonds/salt bridges. Models of protein complexes between human E-cadherin Ec1 domain and InlA/InlB were generated using AlphaFold3. Structures were visualized, analyzed and figures were prepared using PyMol.

To further assess binding specificity, we tested an Ec1 point mutant (Ec1^P16E^) in Pro-16, a key residue in E-cadherin responsible for host specificity towards the internalins of *L. monocytogenes*^28^. This mutation substantially reduced InlB binding compared to wild-type Ec1 (**Figure 4I-J; Figure S7**), confirming the importance of this residue in the interaction. Comparable binding profiles were observed when the same analyses were performed using InlA (**Figure S7-8; Table S1**). Structural modeling further supported these findings, revealing that Ec1 engages the major groove of InlA and InlB through a defined binding interface involving salt-bridge interactions (**Figure 4K-L; Figure S9**). Collectively, these results demonstrate that both InlA and InlB directly and specifically target the Ec1 domain of E-cadherin with high binding affinity.

### InlA and InlB enhance oral epithelial cell migration and invasion

Given that InlA and InlB directly bind the Ec1 domain of E-cadherin, we next examined their interaction with oral epithelial cells and their functional effects on cell motility. IFA showed that FITC-labeled InlA and InlB bound TIGK cells and were localized predominantly to the plasma membrane and partially internalized into the cytoplasm (**Figure 5A, D**). Wound-healing assays showed both InlA and InlB significantly increased TIGK cell migration and wound closure in a time- and dose-dependent manner (**Figure 5B-G; Figure S10**). Transwell assays further showed that InlA and InlB significantly increased TIGK cell invasion (**Fig. 5H, J**) and migration (**Figure 5I, K**). Consistent with these observations, InlB similarly promoted migration and invasion of primary human gingival keratinocytes (**Figure S11**). These pro-migration effects were not observed with a control protein (TDE0471C) from *T. denticola*^46^ which was expressed and purified under identical conditions as InlA/InlB (**Figure S12**) or in RKO^44^, a human colon carcinoma cell line that lacks E-cadherin (**Figure S13**). Additionally, neither InlA nor InlB affected TIGK cell viability or proliferation (**Figure S14A**). Collectively, these findings demonstrate that InlA and InlB specifically enhance oral epithelial cell migration and invasion without altering cell proliferation.

**Figure 5.**
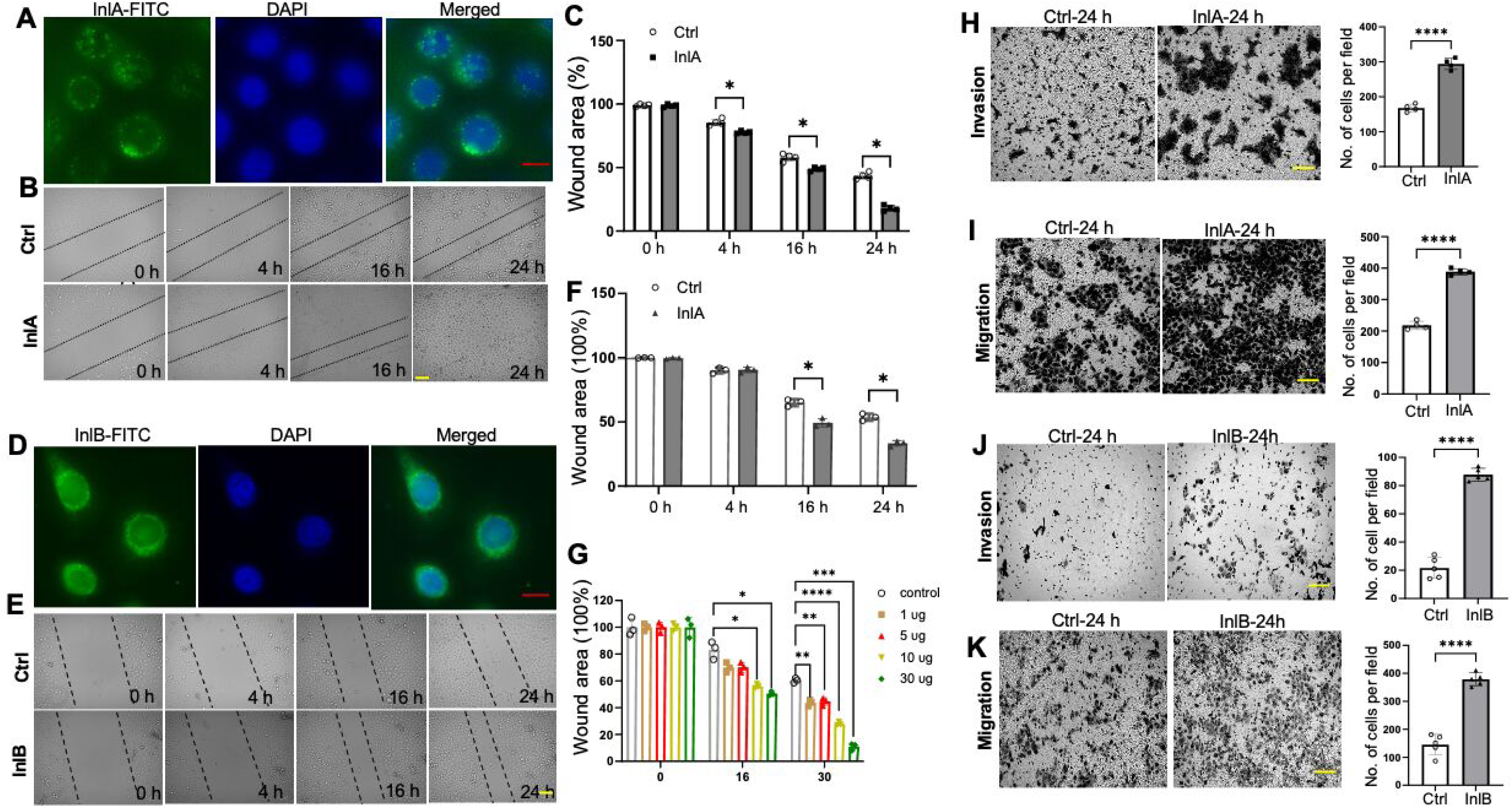
InlA and InlB bind epithelial cells and promote migration and invasion. **(A)** TIGK cells were incubated with FITC-labeled InlA (20 μg/ml, 2 h), and binding was assessed by immunofluorescence microscopy. PBS (Control, Ctrl). Scale bars: 20 μm. **(B-C)** Wound-healing assay evaluating the effects of InlA on TIGK cell migration; wound closure was quantified using ImageJ. Ctrl: PBS. **(D)** TIGK cells were incubated with FITC-labeled InlB (20 μg/ml, 2 h), and binding was analyzed by immunofluorescence. Ctrl: PBS. Scale bars: 20 μm. **(E-F)** Wound-healing assay assessing the effects of InlB on TIGK cell migration; wound closure quantified by ImageJ. Ctrl: PBS. **(G)** Dose-dependent effects of InlB on TIGK cell migration measured by wound-healing assay. **(H-I)** Transwell assays assessing the effects of InlA on TIGK cell migration and invasion; invaded and migrated cells per field were quantified. Ctrl: PBS. **(J-K)** Transwell assays evaluating the effects of InlB on TIGK cell migration and invasion; invaded and migrated cells per field were quantified. Ctrl: PBS.

### InlA and InlB enhance HNSCC epithelial-mesenchymal transition (EMT) and migration

Given that loss of E-cadherin is a hallmark of EMT^20,22^, we examined whether InlA and InlB promote tumor cell motility by inducing EMT-associated signaling pathways. We first assessed E-cadherin expression across eight HNSCC cell lines (SCC1, SCC4, HSC3, SCC9, HN30, SCC47, NU61 and SCC61) and found that only SCC4 lacked E-cadherin expression (**Figure S15**). We then selected HN30 and HSC3 for further investigation and found that treatment of HN30 cells with InlA or InlB significantly enhanced cell migration and invasion, as demonstrated by accelerated wound closure and increased transwell migration and invasion (**Figure 6A-F**), without affecting cell proliferation (**Figure S14B-C**). Consistent with EMT activation, both internalin proteins markedly upregulated expression of the matrix metalloproteinase MMP9 and EMT-associated transcription factors, including ZEB1, Slug, and TWIST, at both the mRNA and protein levels (**Figure 6G-I; Figure S16**). Similar pro-EMT effects were also observed in TIGK and HSC3 cell lines (**Figure S16**). Importantly, knockdown of E-cadherin markedly attenuated InlA- and InlB-induced MMP9 expression (**Figure 6J-L; Figure S17**), indicating that internalin-driven EMT-associated gene induction depends, at least in part, on E-cadherin. Together, these findings demonstrate that InlA and InlB promote HNSCC migration and invasion by promoting EMT in an E-cadherin-dependent manner.

**Figure 6.**
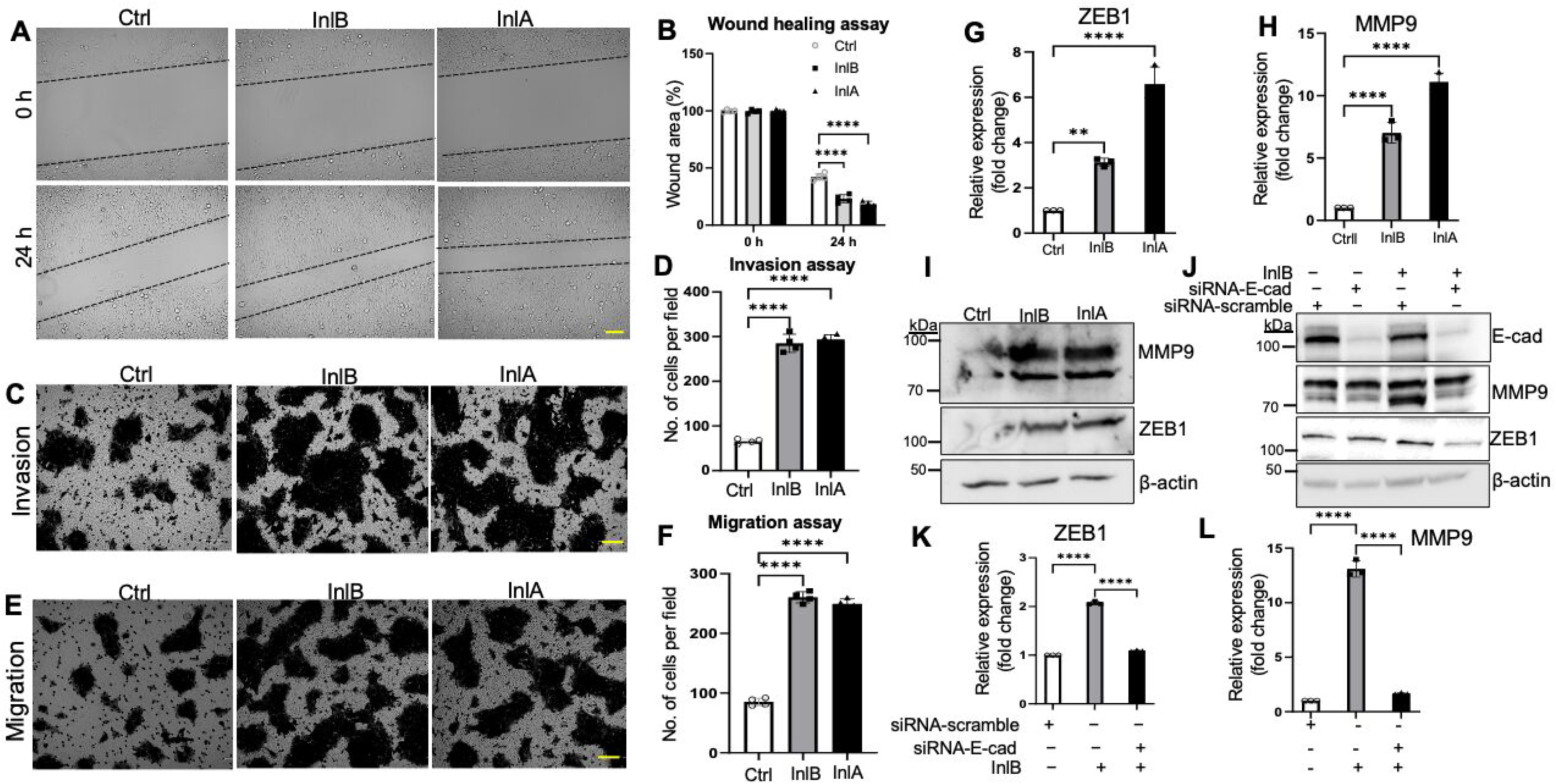
InlB promotes migration and invasion of head and neck cancer cells. **(A-B)** Wound-healing assay assessing HN30 cell migration following InlB treatment (30 μg/ml); representative images at 0 and 24 h (A) and ImageJ quantification of wound area relative to 0 h (B). Ctrl: PBS. **(C-D)**Transwell assays evaluating the effects of InlA or InlB on HN30 cell invasion after overnight treatment; invaded and migrated cells per field were quantified. **(E-F)** Wound-healing assays assessing HN30 cell migration following InlB or InlA treatment; wound area quantified using ImageJ. **(G-H)** qRT-PCR analysis of ZEB1 and MMP9 expression in HN30 cells treated with InlB or InlA for 24 h. **(I)** Immunoblot analysis of ZEB1 and MMP9 protein levels in HN30 cells following InlB or InlA treatment (24 h). **(J)** Immunoblot analysis of ZEB1 and MMP9 in HN3 cells transfected with E-cadherin siRNA and treated with or without InlB or InlA. **(K-L)** qRT-PCR analysis of ZEB1 and MMP9 expression in TIGK and HSC3 cells transfected with E-cadherin siRNA and treated with InlB.

### InlA and InlB induce MMP9 expression via EMT-associated ROCK/JNK/p38 signaling pathways

Previous studies have implicated ROCK (Rho-associated kinase), JNK, p38, and NF-κB pathways in MMP9 regulation during EMT^47,48^. To define the signaling mechanisms underlying InlA- and InlB-induced MMP9 expression, TIGK cells were pretreated with pathway-specific inhibitors prior to internalin stimulation. Inhibition of JNK with SP600125^49^ markedly reduced InlB-induced MMP9 mRNA from ∼20-fold to 4.8-fold (**Figure 7A**), and immunoblotting confirmed strong suppression of MMP9 protein (**Figure 7B**). Likewise, inhibition of NF-κB with cardamonin^50^, ROCK with Y-27632^51^ or p38 with its specific inhibitor IV^52^ substantially attenuated both InlA- and InlB-induced MMP9 protein expression compared to the control groups treated with DMSO (**Figure 7C-D**), indicating contributions from multiple signaling pathways. Consistent with the inhibitor studies, immunoblotting revealed that stimulation with InlA or InlB robustly increased phosphorylation of JNK, p38, NF-κB p65, and the AP-1 component c-Jun, whereas ERK1/2 phosphorylation remained unchanged (**Figure 7E**). Collectively, these findings demonstrate that InlA and InlB selectively bind E-cadherin and activate its downstream an EMT-associated ROCK/JNK/p38 signaling axis that converges on NF-κB and AP-1 transcription factors. This pathway links E-cadherin-targeting internalins to EMT-like transcriptional reprogramming, thereby promoting epithelial cell migration and invasion.

**Figure 7.**
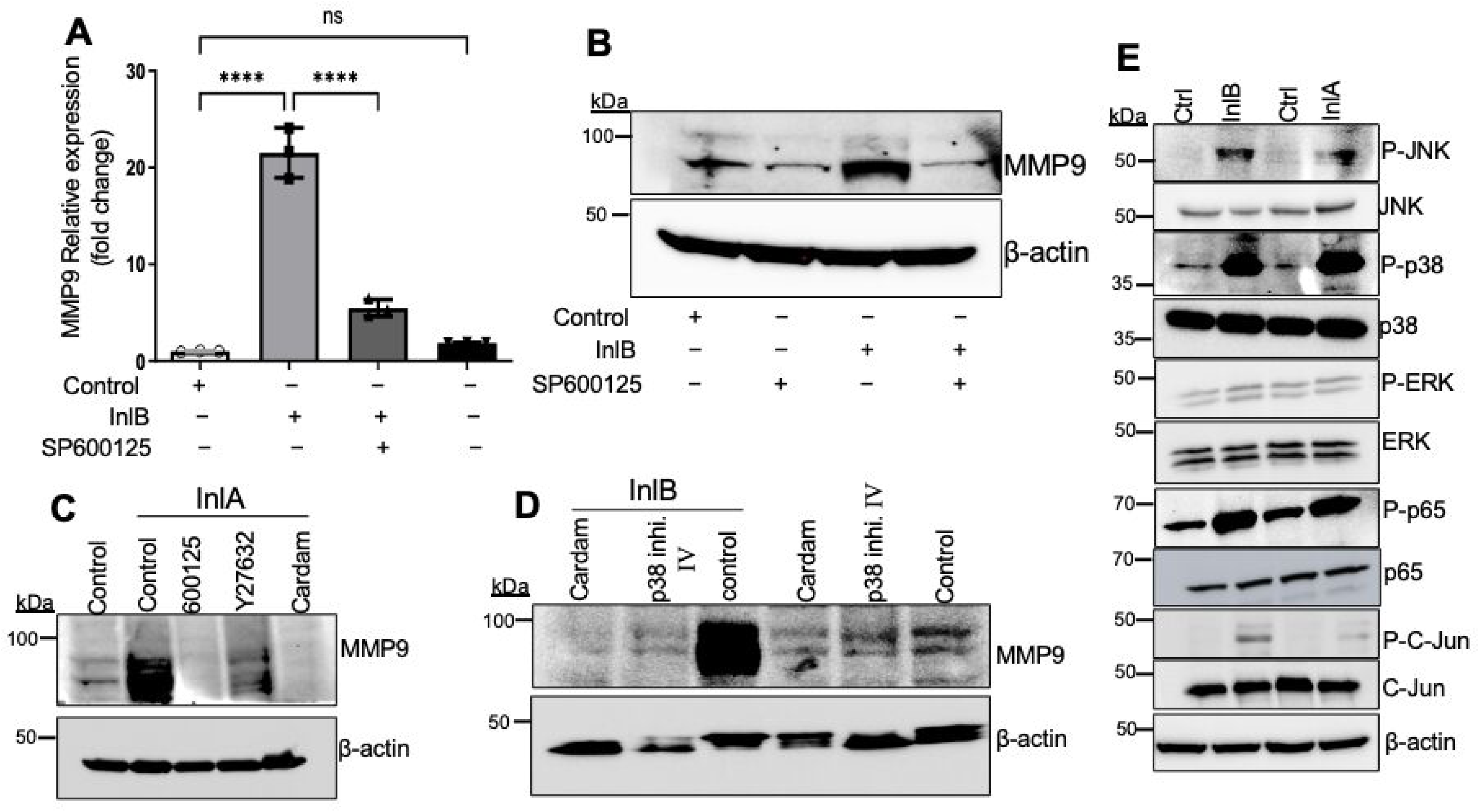
JNK/p38 signaling mediates InlA/InlB-induced MMP9 expression. **(A-B)** InlB-induced MMP9 mRNA and protein expression was suppressed by the JNK inhibitor SP600125. Ctrl: DMSO. **(C-D)** InlA/InlB-induced MMP9 protein levels were inhibited by NF-κB and p38 inhibitors. Ctrl: DMSO. **(E)** Activation of JNK/p38 signaling by InlA and InlB was assessed by immunoblotting for phosphorylated JNK, p38, p65/NF-κB, and c-Jun. Ctrl: PBS.

### InlA and InlB promote β-catenin nuclear translocation via phosphorylation

In epithelial cells, the cytoplasmic tail of E-cadherin forms a dynamic complex with catenins, thereby regulating multiple intracellular signaling pathways^53–55^. Loss or functional impairment of E-cadherin often increases nuclear β-catenin accumulation^22^. To investigate the effects of InlA and InlB on β-catenin, TIGK cells were treated with 20 µg/mL of either protein for 2 hours or overnight, followed by IFA. After 2 hours, β-catenin remained predominantly cytoplasmic, similar to the controls; however, overnight (∼16 h) treatment induced marked β-catenin nuclear translocation (**Figure 8A-B**). Immunoblotting analysis confirmed that nuclear β-catenin levels were significantly elevated, while cytoplasmic levels remained unchanged (**Figure 8C**). To explore its underlying regulatory mechanism, we assessed phosphorylation of β-catenin at Thr41, Ser45, and Tyr654-residues which are implicated in controlling its stability and subcellular distribution^55^. At 16 hours, treatment with InlA or InlB markedly reduced β-catenin phosphorylation at all three sites (**Figure 8D**). Reduced phosphorylation at Thr41 and Ser45 would be expected to inhibit β-catenin degradation and enhance protein stability, thereby promoting its nuclear accumulation. Collectively, these findings suggest that engagement of internalins with E-cadherin promotes β-catenin nuclear enrichment, at least in part by modulating its phosphorylation status and stability.

**Figure 8.**
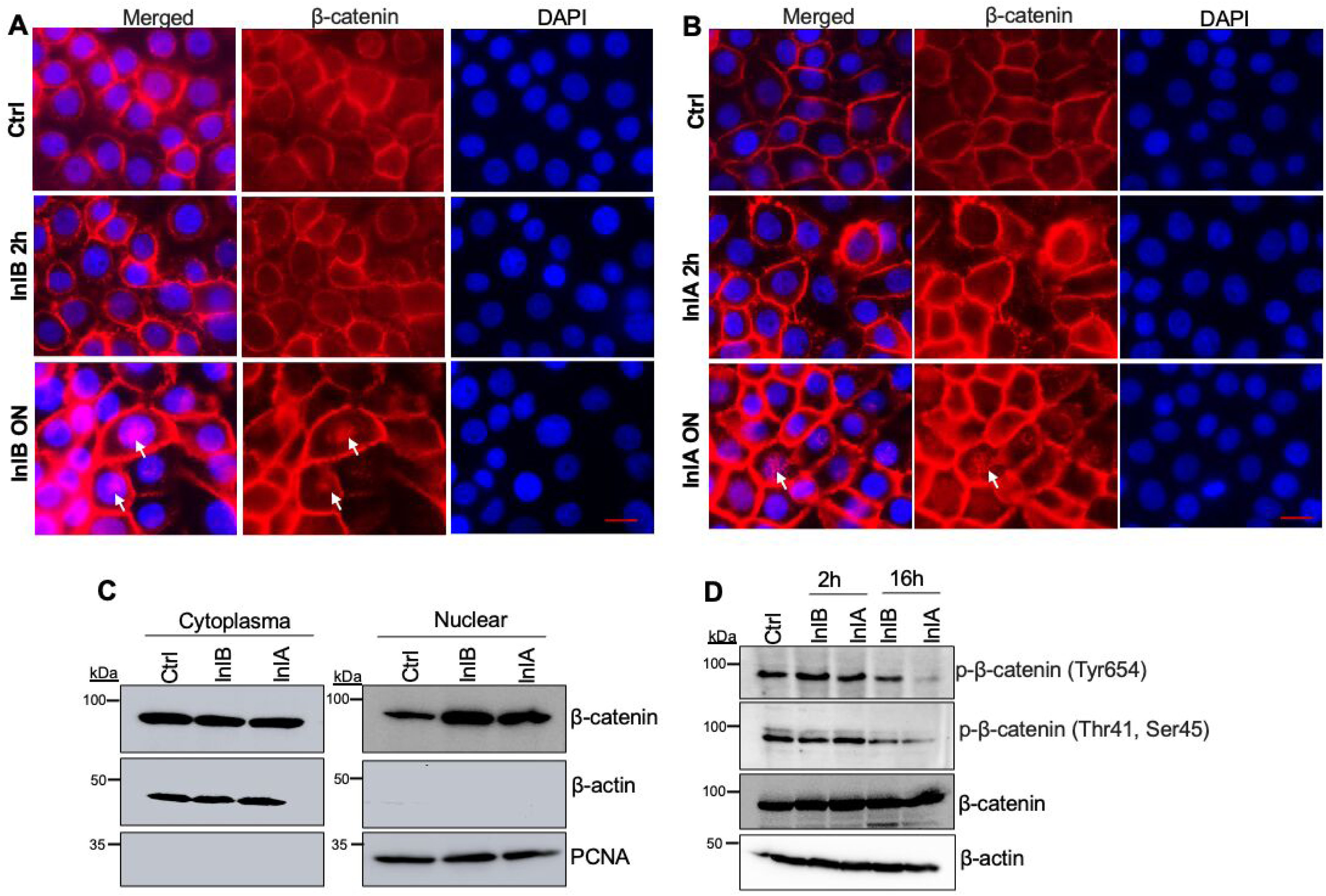
InlA and InlB induce β-catenin phosphorylation and nuclear translocation. **(A-B)** β-catenin localization in TIGK cells treated with InlB (A) or InlA (B) for 2 h or overnight was assessed by immunofluorescence microscopy. Ctrl: PBS. Scale bars: 20 μm. **(C)** Immunoblot analysis of cytoplasmic and nuclear β-catenin levels in TIGK cells following overnight InlB treatment. Ctrl: PBS. **(D)** Immunoblot analysis of β-catenin phosphorylation in TIGK cells treated with InlB for 2 h or overnight. Ctrl: PBS.

### InlA and InlB promote chemoresistance by inhibiting apoptosis

Chronic *P. gingivalis* infection has been linked to tumor multidrug resistance through activation of inflammatory, pro-survival, and drug-efflux pathways^56–58^. To determine whether InlA and InlB contribute to chemoresistance, TIGK, HN30 and HSC3 cells were pretreated with either protein prior to exposure to increasing doses of cisplatin. Cell viability assays revealed that both InlA and InlB significantly (*P < 0.05*) increased cell survival across all tested concentrations (**Fig. 9A-C**). Crystal violet staining confirmed enhanced survival of pretreated cells following the chemotherapy (**Figure 9D**). Using TIGK cells as a surrogate, we showed that InlA and InlB increased cisplatin IC₅₀ from 6.2 µg/mL to 9.3 and 13.2 µg/mL (**Figure 9E**), respectively, indicating reduced sensitivity. Annexin V staining revealed that both proteins protected TIGK and HN30 cells from cisplatin-induced apoptosis (**Figure 9F-H**). Similar effects against 5-FU were observed (**Figure S18**). Mechanistically, InlA and InlB attenuated chemotherapy-induced apoptotic signaling. Immunoblotting analysis revealed that both proteins suppressed cisplatin-induced cleavage of PARP, caspase-3, and caspase-9, reduced expression of the pro-apoptotic protein Bax^59^, and increased levels of the anti-apoptotic factor Bcl-2^60^ (**Figure 9I**). Collectively, these results demonstrate that internalin proteins InlA and InlB confer chemoresistance by suppressing antitumor drug-induced apoptosis, thereby promoting tumor cell survival under chemotherapeutic stress.

**Figure 9.**
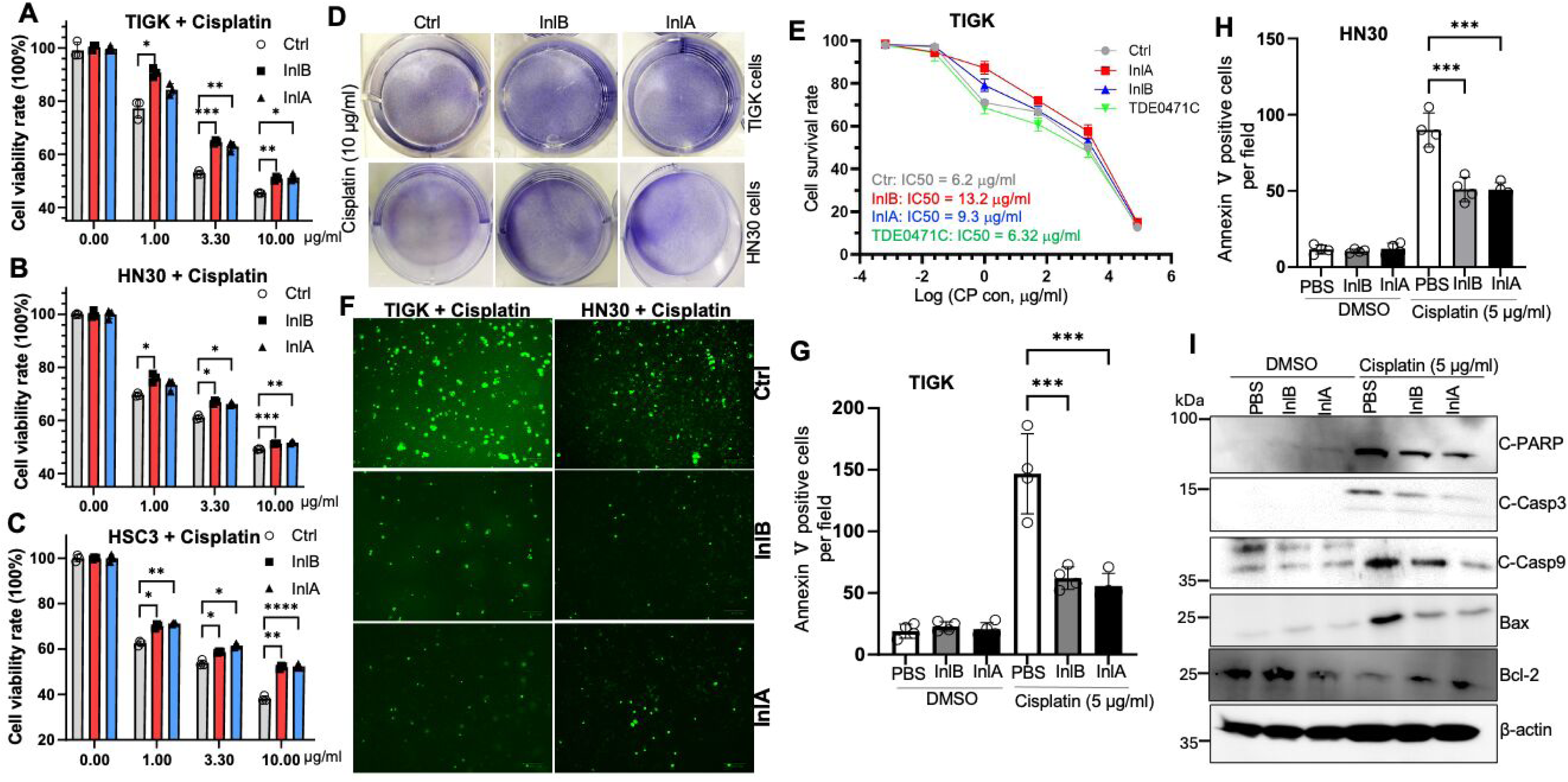
InlA and InlB reduce chemotherapeutic sensitivity by inhibiting apoptosis. **(A-C)** CCK-8 assays determining the viability of TIGK, HN30 and HSC1 cells in cisplatin pretreated with or without InlA/InlB (2 h). Ctrl: PBS. * *P< 0.5*, ** *P< 0.01*, and *** *P< 0.001*. **(D)** Crystal violet staining assessing cell viability following cisplatin treatment in TIGK and HN30 cells pretreated with or without InlA/InlB. Ctrl: PBS. **(E)** CCK-8 assays determining cisplatin IC₅₀ in TIGK cells pretreated with or without InlA/InlB (2 h). Ctrl: PBS. **(F-H)** Annexin V staining detecting cisplatin-induced apoptotic cells of TIGK and HN30 pretreated with or without InlA/InlB (2 h). Ctrl: PBS. ** *P< 0.01*, and *** *P< 0.001*. **(I)** Immunoblot analysis of apoptosis-related proteins (cleaved caspase-3, cleaved caspase-9, cleaved PARP, Bcl-2, and Bax) in cells treated with cisplatin with or without InlA/InlB pretreatment. Ctrl: PBS.

### InlA and InlB promote HNSCC lung metastasis *in vivo*

To assess the *in vivo* effects of InlA and InlB on tumor metastasis, HN30-luciferase cells were pretreated with either protein for 2 hours and injected into NSG mice via the tail vein. Bioluminescence imaging revealed that pretreated cells exhibited markedly increased lung metastasis compared with controls (**Figure 10A-B**). Five weeks post-injection, ex *vivo* IVIS imaging confirmed a substantial increase in total photon flux in lungs from InlA- or InlB-treated tumor cells (**Figure 10C-D**). Gross examination showed more visible tumor nodules and higher lung weights in these groups (**Figure 10E-F**). Consistently, histological analysis of H&E-stained lung sections showed a dramatic increase in metastatic nodules (InlA: 81.0 nodules/lung; InlB: 80.2 nodules/lung; control: 14.2 nodules/lung) and greater tumor burden, as reflected by the percentage of lung area occupied by tumor tissue (InlA: 27.8%; InlB: 30.8%; control: 17.3%) (**Figure 10G-I**). Together, these data demonstrate that InlA and InlB significantly enhance HNSCC lung metastasis *in vivo*, supporting a pro-metastatic role of *P. gingivalis* internalin proteins.

**Figure 10.**
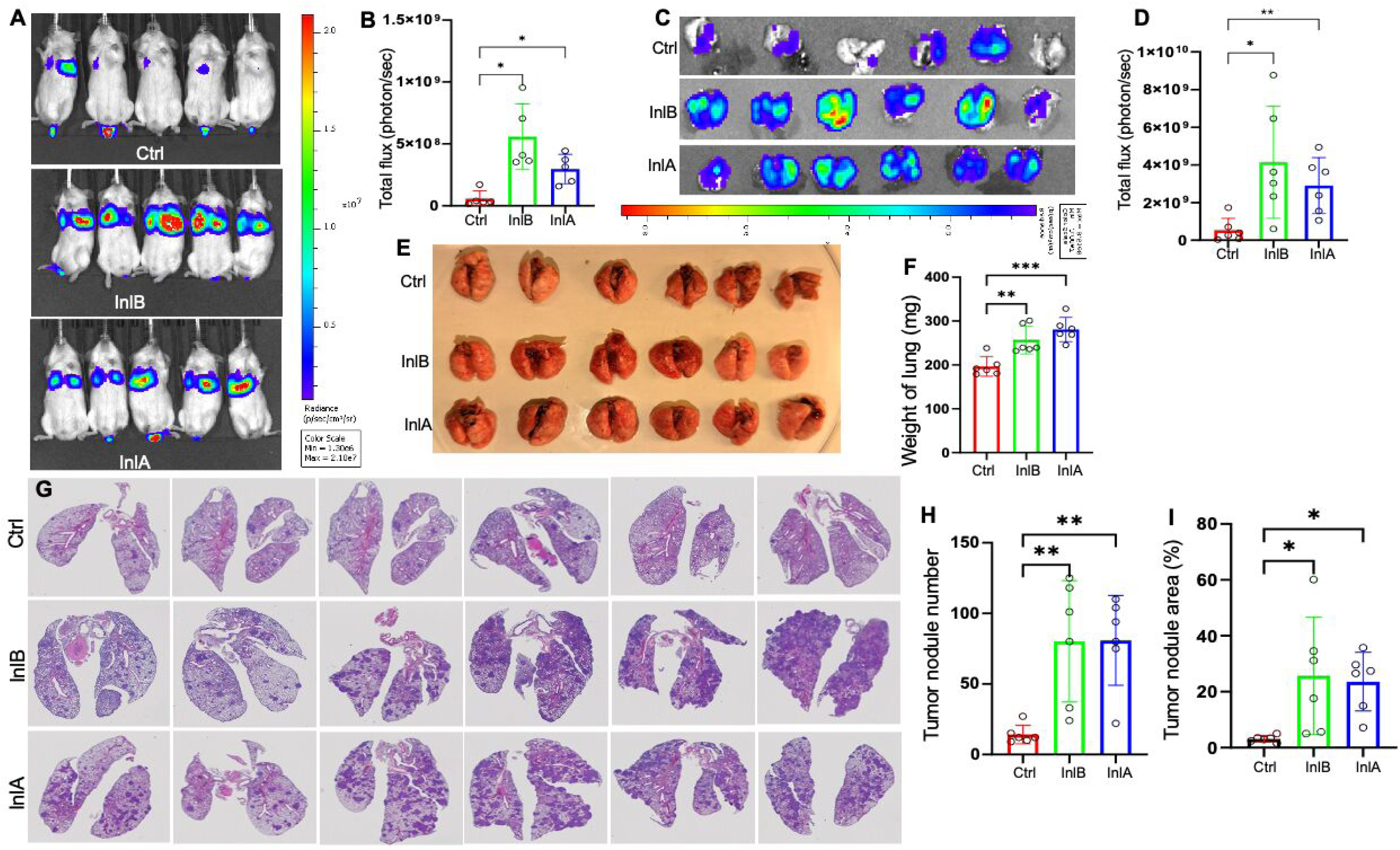
InlA and InlB promote tumor metastasis in vivo. **(A-B)** Whole-body bioluminescence of mice was imaged using the IVIS system, and luciferase flux was quantified. Ctrl: PBS. **(C-D)** Bioluminescence in the lungs of each group was measured by IVIS, and total luciferase flux was quantified. Ctrl: PBS. **(E-F)** Lungs were collected, and average lung weight per group was measured and quantified. Ctrl: PBS. **(G)** Hematoxylin and eosin (H&E) staining showing lung tumor nodules. Ctrl: PBS. **(H)** Quantification of tumor nodule number per lung section across groups. Ctrl: PBS. **(I)** Tumor nodule area relative to total lung area was measured using ImageJ. Ctrl: PBS.

### Detection of InlB in HNSCC patient tissues

A tissue microarray was used to assess the presence of *P. gingivalis* internalin proteins in HNSCC specimens. The array comprised 48 samples (24 tumors and 24 adjacent tissues) from 12 patients, representing gingiva, larynx, palate, salivary gland, tongue, and maxillary tissues. Immunohistochemical (IHC) analysis using an anti-InlB antibody identified 14 positive cases (29%), including 4 with strong staining (3+, H-score ≥150) and 10 with moderate staining (2+, H-score ≥80) (**Figure 11**). These results demonstrate that InlB is present within tumor tissues, consistent with its secretion and exposure in the tumor microenvironment, and align with emerging evidence implicating intratumoral bacteria in HNSCC and related malignancies.

**Figure 11.**
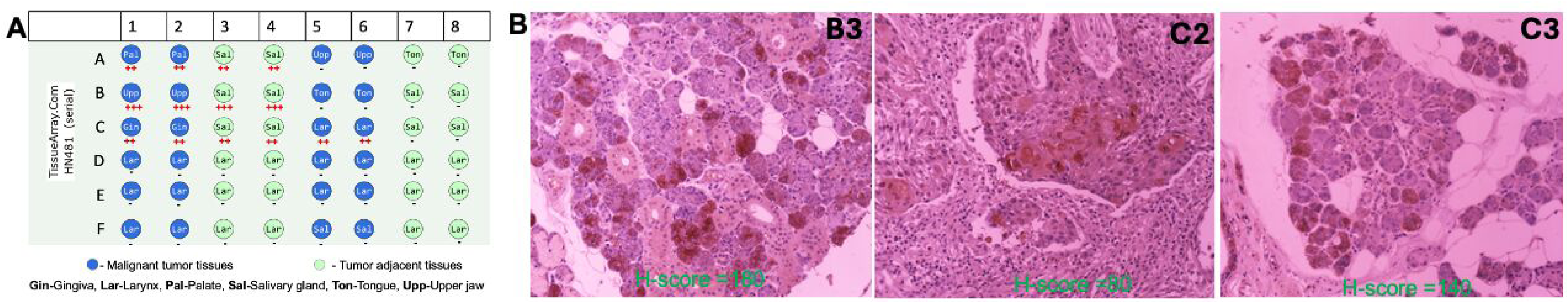
Detection of InlB in HNSCC patient tissues using immunohistochemistry (IHC). **(A)** Diagram illustrating the tissue microarray containing different tissues from HNSCC patients. **(B)** Representative IHC positive images. Sections were incubated overnight at 4 °C with a primary antibody against InlB (generated in-house; 1:300 dilution). Signals were detected using a biotinylated secondary antibody and streptavidin-HRP, followed by visualization with DAB and counterstained with Mayer’s hematoxylin. Brown DAB chromogen indicates bacterial antigen positivity.

## DISCUSSION

HNSCC is among the most common cancers worldwide^1^. Clinical and epidemiological studies link periodontal disease and *P. gingivalis* infection with an increased risk of HNSCC^5^, and growing experimental evidence demonstrates that this oral pathogen promotes tumorigenesis by sustaining inflammation, suppressing apoptosis, and enhancing EMT and chemoresistance^16,17,58,61^. However, the specific bacterial effectors that mediate these oncogenic effects have remained poorly defined. Here, we identify InlA and InlB, two internalin-like proteins, as central drivers of *P. gingivalis* invasion and tumor progression. Secreted via T9SS, InlA and InlB bind specifically to human E-cadherin and promote bacterial entry into gingival epithelial cells (**Figures 2-4**).

Beyond facilitating cell invasion, engagement of E-cadherin by InlA and InlB profoundly alters host signaling. E-cadherin normally functions as a tumor suppressor by sequestering β-catenin at AJs and restraining EMT^21,30,62^. InlA/InlB binding disrupts this regulatory role, leading to β-catenin release and nuclear translocation and activation of EMT-associated transcriptional regulators (**Figures 6-8**). In parallel, InlA- and InlB-mediated activation of JNK/p38 MAPK pathways stimulates c-Jun- and NF-κB-dependent MMP9 expression (**Figures 6-7**) and leads to activation of pro-survival signaling (**Figure 9**). Collectively, these coordinated effects enhance HNSCC cell migration (**Figure 5**), chemoresistance (**Figure 9**), and metastatic potential *in vivo* (**Figure 10**). A working model (**Figure 12**) is proposed to illustrate the mechanism by which InlA and InlB convert E-cadherin from a static adhesion molecule into a signaling hub that drives EMT and therapeutic resistance, providing a direct mechanistic link between chronic oral infection and tumor progression.

**Figure 12.**
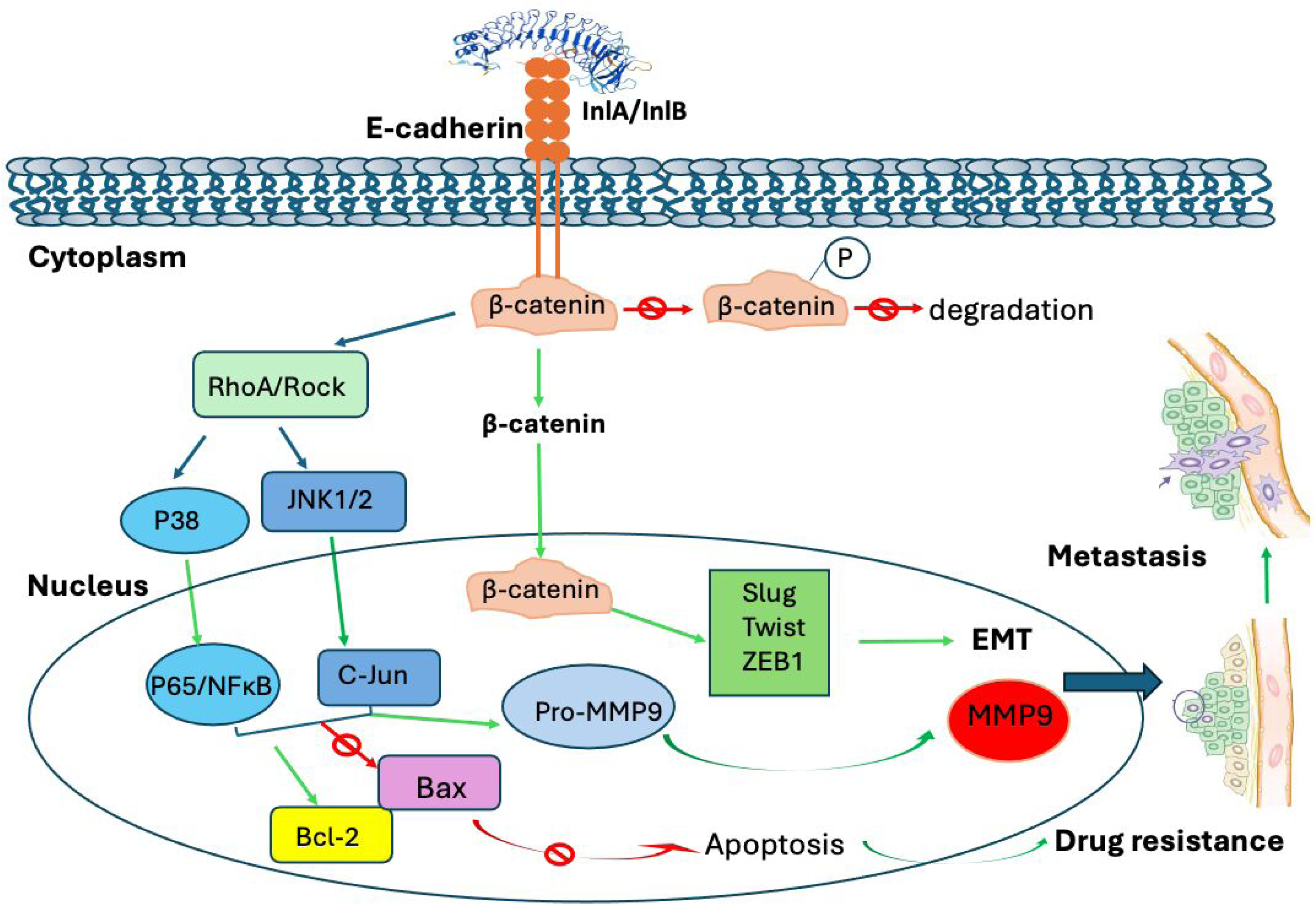
Mechanistic model of *P. gingivalis* InlA/InlB in metastasis and drug resistance. InlA and InlB bind E-cadherin, triggering β-catenin stabilization and nuclear translocation, which induces EMT transcription factors (Slug, Twist, ZEB1) and MMP9, promoting metastasis. Concurrently, internalin-E-cadherin engagement activates downstream ROCK, p38 and JNK MAPK signaling, attenuating cisplatin-induced apoptosis through upregulation of Bcl-2 and suppression of Bax, ultimately conferring chemoresistance. These coordinated pathways mechanistically link *P. gingivalis* infection to cancer progression and therapeutic resistance. Green arrows denote activation; red blocked arrows denote inhibition.

Bacterial internalins are classically regarded as invasion factors that promote host cell entry through receptor engagement^25,27,29^. Canonical internalins from *L. monocytogenes* bind epithelial receptors including E-cadherin and the tyrosine kinase Met to induce localized cytoskeletal rearrangements^25,27,63^. Although some internalins can transiently modulate inflammatory or survival pathways^29,64–66^, their capacity to drive sustained, disease-relevant reprogramming of epithelial signaling, particularly in the context of cancer, has remained largely unexplored. Our findings expand this conceptual framework by demonstrating that *P. gingivalis* InlA and InlB couple bacterial invasion to oncogenic signaling. Although they functionally resemble canonical internalins in targeting E-cadherin, their secretion via the T9SS enables receptor engagement independent of direct bacterial contact, allowing sustained modulation of host signaling in chronically infected tissues such as the gingiva and tonsils. In contrast, *L. monocytogenes* internalins, including InlA and InlB, are typically anchored to the bacterial cell wall through C-terminal “LPXTG” or “GW motifs”^29,67^. Moreover, *P. gingivalis* internalins display limited sequence similarity and distinct domain architectures compared with their *Listeria* counterparts; for example, *L. monocytogenes* InlA and InlB contain characteristic Ig-like interrepeat domains^27,41^. Together, these differences highlight the evolutionary and functional divergence of internalin-like proteins across bacterial species.

Given its critical role in maintaining AJs integrity^20,21,30^, multiple bacterial pathogens have evolved to exploit E-cadherin receptor. For example, *Fusobacterium nucleatum* FadA destabilizes E-cadherin to activate β-catenin-dependent proliferation^24^, *Bacteroides fragilis* enterotoxin cleaves E-cadherin to promote inflammation and growth^68^, and *Helicobacter pylori* CagA disrupts AJs to sustain intracellular oncogenic signaling^69^. In contrast, *P. gingivalis* InlA and InlB employ a distinct strategy: rather than degrading or destabilizing E-cadherin, they reprogram it into a persistent signaling platform. Through T9SS-mediated secretion, InlA and InlB engage E-cadherin beyond sites of direct bacterial contact, linking invasion to coordinated activation of β-catenin and ROCK-JNK/p38 MAPK signaling pathways. This integrated signaling output drives EMT, MMP9 expression, and chemoresistance, thereby promoting tumor progression. To our knowledge, this represents the first example of bacterial internalins converting a tumor suppressor into a pro-metastatic signaling hub, establishing a new paradigm for how oral pathogens directly contribute to cancer progression through defined molecular effectors.

To place InlA and InlB in an evolutionary context, we searched all sequenced *P. gingivalis* strains and found conserved homologs in each genome. Expanding this analysis identified 385 related proteins predominantly within the Cytophaga-Flavobacterium-Bacteroides (CFB) phylum, which possesses the T9SS secretion apparatus^70^, with occasional representation in Firmicutes (**Figure S19**). Phylogenetic analysis revealed that these proteins cluster with a T9SS type A sorting domain-containing protein from *Porphyromonas* species, suggesting that this family of proteins originated from ancestral T9SS-secreted proteins. The presence of Firmicutes sequences within otherwise CFB-dominated clades may reflect horizontal gene transfer or retention from a common ancestor. Together, these findings position InlA and InlB within a broader family of T9SS-associated internalin-like proteins and suggest that coupling invasion functions with secreted, long-range receptor engagement may represent a conserved strategy among oral and gut-associated CFB bacteria.

In conclusion, this work demonstrates that *P. gingivalis* internalins act as potent modulators of epithelial signaling, cancer cell plasticity, and therapeutic resistance. By defining a mechanistic link between bacterial E-cadherin engagement and EMT-associated signaling, our findings reveal how chronic infection can function as a non-genetic driver of cancer progression and underscore the broader impact of host-microbe interactions on tumor biology.

### Limitations of this study

In this study, we examined the roles of InlA and InlB in tumor cell migration, metastasis, and chemoresistance using purified proteins rather than intact *P. gingivalis* cells, thereby minimizing confounding contributions from additional virulence determinants such as gingipains and lipopolysaccharide. Although this reductionist approach enabled direct evaluation of protein-specific activities, it does not fully recapitulate the complexity of host-pathogen interactions during infection. In vivo metastasis assays were performed using tail vein injection models, which primarily assess late-stage metastatic colonization rather than primary tumor growth and spontaneous dissemination. Future studies will define the structural basis of InlA/InlB engagement with the Ec1 domain of E-cadherin and evaluate their roles in tumor progression and microenvironmental remodeling using orthotopic xenograft models.

## ACKNOWLEDGEMENT

This project is supported by DE023080, DE030667 and DE034063 to C. Li and DE033686 to X. Wang. We thank Dr. Yi-ping Han for providing E-cadherin constructs and RKO cell line. We also thank Dr. Zhao-Qing Luo for providing plasmids for yeast toxicity assays, Dr. Janina Lewis for providing pG108 shuttle vector and Dr. Xinyan Pei for her assistance with SPR. Services and products in support of the research project were generated by the Virginia Commonwealth University Flow Cytometry Shared Resource, supported, in part, with funding from NIH-NCI Cancer Center Support Grant P30 CA016059.

## Experimental Procedures

### Reagents and antibodies

Recombinant human E-cadherin protein was purchased from Fisher Scientific (cat. no. 50-101-7679). JNK inhibitor SP600125 (S5567), NF-κB inhibitor cardamonin (cat. no. 481406), and p38 MAP kinase inhibitor IV (cat. no. SML0543) were obtained from Millipore-Sigma. The ROCK inhibitor Y-27632 (cat. no. 1254) was purchased from Tocris Bioscience. Primary antibodies used in this study include anti-His (Invitrogen, cat. no. MA1-21315); E-cadherin (R&D Systems, cat. no. AF648); β-catenin (R&D Systems, cat. no. MAB1329); AKT (Cell Signaling Technology, cat. no. 9272); phospho-AKT (Ser473; cat. no. 9271); p38 (cat. no. 9212); phospho-p38 (Thr180/Tyr182; cat. no. 9211); JNK (cat. no. 9252); phospho-JNK (Thr183/Tyr185; cat. no. 9251); NF-κB p65 (cat. no. 8242); phospho-NF-κB p65 (Ser536; cat. no. 3033); c-Jun (cat. no. 9165); phospho-c-Jun (Ser63; cat. no. 9164); ERK1/2 (cat. no. 9102); phospho-ERK1/2 (Thr202/Tyr204; cat. no. 4370); and β-actin (cat. no. 3700) (all from Cell Signaling Technology unless otherwise indicated). Antibodies against MMP9 (cat. no. MA5-15886), SNAIL (cat. no. 14-9859-82), SLUG (cat. no. PA5-20289), TWIST (cat. no. MA5-17195), and ZEB1 (cat. no. 14-9741-82) were purchased from Thermo Fisher Scientific.

### Bacterial strains and growth conditions

All *Porphyromonas gingivalis* strains were cultured in trypticase soy broth (TSB) supplemented with yeast extract (5 µg/mL), hemin (5 µg/mL), and vitamin K (1 µg/mL) at 37 °C in an AS-500 anaerobic chamber (Biolog, Hayward, CA) at atmosphere of 85% N₂, 10% H₂, and 5% CO₂, as previously described^71,72^. Solid cultures were maintained on trypticase soy agar containing 5% defibrinated sheep blood and the same supplements. Where indicated, clindamycin (1 µg/mL) and/or tetracycline (1 µg/mL) were added. *Escherichia coli* NEB5α was used for DNA cloning, and BL21-Star (DE3) was used for recombinant protein expression. ClearColi BL21(DE3) (Biosearch Technologies) was used for preparation of endotoxin free recombinant proteins. *E. coli* strains were cultured in lysogeny broth (LB) supplemented with appropriate antibiotics unless otherwise noted.

### Construction and validation of *P. gingivalis* mutants

Genes encoding InlA (PG0350), InlB (PG1374), and PorT (PG0751) were inactivated in *P. gingivalis* W83^73^ by insertion of, or in-frame replacement with, an erythromycin resistance cassette (*ermF*), as previously described^71,72^. Briefly, 280-380 bp fragments corresponding to the 5′ and 3′ flanking regions of each target gene were PCR-amplified and fused to *ermF* by two-step PCR. The resulting constructs were cloned into pJET1.2, sequence verified, and electroporated (∼1 μg) into W83 competent cells. Transformants were recovered under anaerobic conditions and selected on TSA blood agar containing clindamycin (1 μg/mL). Deletion mutants (*ΔinlA*, *ΔinlB*, and *ΔporT*) were confirmed by PCR. For complementation, full-length *InlA* or *InlB* gene, including their native promoters, were cloned into the pG108 shuttle vector^74^ by Gibson assembly, sequence verified, and electroporated (∼5 μg) into the corresponding mutant strains. Complemented strains (*cΔinlA* and *cΔinlB*) were selected on TSB blood agar containing clindamycin and tetracycline (1 μg/mL each), screened by PCR, and confirmed by immunoblotting using InlA- or InlB-specific antibodies. Growth of *P. gingivalis* W83 and mutant strains was monitored by measuring optical density at 600 nm (OD600) using a spectrophotometer (Thermo Fisher Scientific). Primers used in this study are listed in Supplementary Table 1.

### Production of recombinant proteins and antibodies and site-directed mutagenesis

The nucleotide sequence encoding InlA without its N-terminal signal peptide (26-484 residues) was codon-optimized and subcloned into the pET101 expression vector. The resulting construct was transformed into *E. coli* BL21-Star (DE3) or ClearColi BL21(DE3) for protein expression using LB media with IPTG induction with shaking at 16°C overnight. The nucleotide sequence encoding InlB lacking its N-terminal signal peptides (23-428 residues) was PCR-amplified from the W83 strain, cloned into pJET1.2, sequence verified, and subcloned into the pQE80 expression vector. The resulting constructs were transformed into *E. coli* BL21-Star (DE3) or ClearColi BL21(DE3) for protein expression in auto-induction medium with shaking overnight at 37°C.

Recombinant InlA and InlB proteins were purified under native conditions using a HisTrap HP Ni-NTA column on an NGC FPLC system (Bio-Rad), followed by size-exclusion chromatography (SEC) on a Superdex 200 Increase 10/300 GL column (Cytiva Life Sciences) for further purification. For E-cadherin recombinant proteins, nucleotide sequences encoding the ectodomains (Ec1-5, Ec2-5, and Ec1) were PCR-amplified from pCDNA-hE-cadherin (a gift from Dr. Yiping Han)^24^, cloned into pJET1.2, sequence verified, and subcloned into the pGEX-6p-1 expression vector carrying an N-terminal GST tag. In addition, Ec1^P16E^, a point mutant in Ec1 proline-16 (P16E), was created using a Q5 site-directed mutagenesis kit (NEB). The resulting constructs were transformed into BL21-Star (DE3) for protein expression induced with 1 mM IPTG at 37°C for 4 h (Ec1-5 and Ec2-5) or at 16°C overnight (Ec1 and Ec1^P16E^). The resulting GST fusion recombinant proteins were purified using glutathione Superflow agarose resin according to the manufacturer’s instructions. Purified InlA, InlB, and E-cadherin proteins were pooled, dialyzed into dPBS, concentrated using Spin-X UF centrifugal concentrators (5 or 10 kDa cutoff; Corning), and quantified by bicinchoninic acid (BCA) assay (Pierce). Antisera against InlA and InlB were raised in rabbits at Fee-for-service (General Bioscience Corporation) following a standard immunization protocol.

### Yeast manipulation

*Saccharomyces cerevisiae* strain W303 was grown at 30°C in YPD medium or in synthetic media with appropriate amino acid dropout supplemented with either 2% glucose or galactose as the sole carbon source. Yeast transformation was performed using the standard lithium acetate protocol. For yeast lethality assay, DNA fragments encoding InlA (26-484 residues) and InlB (24-428 residues) were cloned into pYES2/NTA plasmid (a gift from Z. Q. Luo from Purdue University) that carries the galactose inducible Pgal1 promoter. After 3 days incubation at 30°C, the resultant transformed clones were confirmed by immunoblotting analysis. For yeast lethality assay, overnight cultures of yeast transformants were serially diluted and spotted on appropriate amino acid dropout supplemented with either 2% glucose or galactose plates. The primers for constructing the plasmids are listed in **Table S2**.

### Transient expression of InlA and InlB in HEK293T cells

The DNA fragments encoding InlA and InlB were codon optimized, synthesized by GenScript Biotech (Piscataway, NJ), and cloned into pcDNA6.2 C-EmGFP-GW TOPO expression vector (Invitrogen, Waltham, MA). For transfection, a total of 6.25 × 10⁵ HEK293T cells were mixed with 4 µg of transfection vector and 16 µl of FuGENE transfection reagent (Promega, Madison, WI) in 400 µl of Opti-MEM medium (Gibco, Waltham, MA). The mixture was then seeded into 24-well cell culture plates containing German glass coverslips (Thermo Fisher Scientific, Waltham, MA) and cultured at 37 °C with 5% CO₂ for 24 h. After incubation, the culture medium was replaced with DMEM containing 10% FBS, and the cells were cultured for an additional 24 h. The cells were then collected and subjected to immunofluorescence imaging. Primers used here are listed in **Table S2.**

### Immunofluorescence staining

Immunofluorescence staining was performed as previously described^75^. Briefly, TIGK^76^, HSC3^77^, or HN30 cells^78^ (2.5 × 10^5^ cells per well) were seeded into 12-well plates and cultured overnight. Cells were treated with FITC-conjugated InlA/InlB (20 µg/ml) for 2 h, fixed with 3.7% paraformaldehyde for 30 min at room temperature (RT), and permeabilized with 0.1% Triton X-100 for 10 min. After washing with PBS, cells were blocked with 5% FBS in PBS for 1 h and incubated with anti-E-cadherin primary antibodies overnight at 4°C. Cells were then washed and incubated with Alexa Fluor 594-conjugated secondary antibodies at RT for 1 h. Nuclei were counterstained with DAPI, and coverslips were mounted using antifade mounting medium. Colocalization of FITC-labeled InlA/InlB and E-cadherin was analyzed using a Zeiss fluorescence microscope equipped with a ×63 oil immersion objective (numerical aperture 1.4).

### ELISA-based binding assay

An ELISA was performed to assess the interaction between InlA/InlB and E-cadherin. Purified His-tagged InlA/InlB proteins (1 µg in 100 µL PBS, pH 7.4) was immobilized in high-binding 96-well microtiter plates (Greiner Bio-One) by overnight incubation at 4°C. Wells were washed with PBS and blocked with 2% bovine serum albumin (BSA) in PBS at RT for 1 hour, followed by incubation with recombinant human E-cadherin (Fisher Scientific, cat. no. 50-101-7679) at the indicated concentrations at RT for 2 h. After washing, bound E-cadherin was detected using an anti-E-cadherin primary antibody, followed by horseradish peroxidase (HRP)-conjugated sheep anti-goat IgG (H+L) secondary antibody (Invitrogen). Binding was quantified by measuring absorbance at 450 nm following enzymatic conversion of 3,3′,5,5′-tetramethylbenzidine (TMB; Thermo Fisher Scientific).

### Surface Plasmon Resonance (SPR) analysis

SPR experiments were performed using a Reichert SR7500DC system, and data were analyzed with TraceDrawer software (v1.8.1), as previously described^75^. Recombinant InlA/InlB proteins (ligands) were immobilized onto planar polyethylene glycol/carboxyl or CM5 carboxymethyl dextran sensor chips via standard amine coupling using N-hydroxy-succinimide (NHS) and N-ethyl-3-(3-dimethylaminopropyl)-carbodiimide (EDC). Chips were pre-conditioned with running buffer (dPBS with 0.01-0.1% Tween-20 and 0.1-0.25% BSA), activated with freshly prepared EDC/NHS, and exposed to 100-500 μg/mL ligand in dPBS, with immobilization repeated three times. Surfaces were then blocked with 1 M ethylenediamine and 1 M ethanolamine. For binding assays, purified E-cadherin ectodomains (analytes) were injected at 500 nM to 8-10 μM in the respective running buffers. Following association, buffer alone was applied to monitor dissociation, and surfaces were regenerated between injections with 10 mM glycine (pH 2.0). Binding kinetics were analyzed by globally fitting data to multiple binding models, and EC_50_ values were calculated using an affinity model.

### Cell invasion and flow cytometry assays

Human telomerase-immortalized gingival keratinocytes (TIGKs)^76^ were maintained in keratinocyte serum-free growth medium (KGM; Lifeline Cell Technology) at 37°C in 5% CO₂. *P. gingivalis* invasion assays were performed as previously descried^39,79^ with some modifications. In brief, TIGKs were seeded at 5 × 10⁵ cells/well in 6-well plates and grown to confluency. Overnight *P. gingivalis* cultures were grown to mid-log phase (OD_600_ 0.5-0.7), washed with PBS, and labeled with 5 μM BCECF-AM (ThermoFisher Scientific) for 30 min at 37°C under anaerobic conditions in the dark, followed by PBS washes to remove residual dye. TIGKs were equilibrated with anaerobic KGM and infected with BCECF-AM-labeled *Pg* cells at an MOI of 100 or 200 for 1 to 2 h at 37°C under anaerobic conditions. After infection, TIGK cells were washed, detached with trypsin-EDTA, pelleted, resuspended in KGM, and analyzed by flow cytometry using a FACSymphony A1 (BD Biosciences). To distinguish internalized bacteria from surface-bound bacteria, cells were quenched with trypan blue prior to analysis. Negative controls include *P. gingivalis* cells incubated with DMSO (vehicle for BCECF-AM) and noninfected TIGKs for gating. Percent invasion was calculated as the percentage of invaded cells among 10,000 events, and data are presented as the mean ± SEM from three independent experiments performed in duplicate.

### Cell lines, culture conditions, and treatments

The human tongue squamous cell carcinoma line HSC3 was purchased from Millipore-Sigma (cat.# SCC193). The HN30-Luciferase (HN30-Luc), SCC61/NU61 and SCC47 cell lines were provided by Dr. Iain Morgan (Philips Research Institute, VCU). Primary gingiva keratinocytes (PCS-200-014), SCC9 (CRL-1629) and SCC4 (CRL-1624) cell lines were from ATCC. The human RKO cell line was obtained from Dr. Yiping Han (Columbia University). MOCI cell line was obtained from Dr. Xiang-yang Wang. SCC1 cells were provided by Dr. Jiong Li (School of Pharmacy, VCU). HSC3 and all SCC cell lines were cultured in DMEM supplemented with 10% fetal bovine serum (FBS) and 1% penicillin-streptomycin at 37°C in a humidified atmosphere containing 5% CO₂. HN30-Luc cells were maintained in DMEM supplemented with 10% FBS, 1% penicillin-streptomycin, and blasticidin (10 µg/mL).

For gene expression analyses, cells were treated with purified recombinant InlA or InlB proteins (20-30 µg/ml) for the indicated times and harvested for analysis of MMP2, MMP9, and EMT-related markers (Snail1, Slug, ZEB1, and TWIST) at the mRNA by qRT-PCR and protein levels by immunoblotting. For signaling pathway activation assays, cells were treated with recombinant InlA or InlB (20-30 µg/ml) for 15 min and harvested for immunoblot analysis of phosphorylated p38, JNK, AKT, c-Jun, p65, and ERK, along with their corresponding total protein levels.

For pathway inhibition studies, TIGK, HN30, or HSC3 cells were pretreated for 2 h with the ROCK inhibitor Y-27632 (10 µM), the JNK inhibitor SP600125 (50 µM), the NF-κB inhibitor cardamonin (20 µM), or p38 MAP kinase inhibitor IV (20 µM), followed by treatment with recombinant InlA or InlB proteins (20-30 µg/ml) for 24 h. Cells were then harvested for RNA or protein isolation to assess the expression of MMP2, MMP9, and EMT-related markers by qRT-PCR or immunoblotting.

### Colony formation, Transwell invasion/migration, and wound-scratch assays

For colony formation assays, 2,000 TIGK, HSC3, or HN30 cells were seeded into 6-well plates and treated with PBS (control) or recombinant InlA/InlB proteins (20 µg/ml). Cells were cultured for 7 days, fixed with methanol for 30 min, and stained with 0.5% crystal violet for 10 min. Colonies containing more than 50 cells were counted under a light microscope.

For Transwell invasion assays^80^, 2 × 10⁵ TIGK, HSC3, or HN30 cells suspended in serum-free medium were seeded into the upper chambers of 24-well Transwell inserts coated with Matrigel and treated with or without recombinant InlA or InlB proteins (20 µg/ml). DMEM containing 10% FBS (600 µL) was added to the lower chamber as a chemoattractant. After overnight incubation, non-invading cells on the upper surface were removed with a cotton-tipped applicator. Invaded cells on the lower surface were fixed with ice-cold methanol for 10 min, stained with 0.5% crystal violet for 10 min, washed with water, air-dried, and imaged by light microscopy. Migration assays were performed using the same protocol, except inserts were not coated with Matrigel.

For wound-scratch assays^81^, 5 × 10⁵ TIGK, HSC3, or HN30 cells were seeded into 6-well plates and cultured overnight to confluence. Cells were treated with recombinant InlA or InlB at the indicated concentrations (1∼ 30 µg/ml), and a linear scratch was generated using a 200-µL pipette tip. Images were captured at 0, 6, 16, and 24 h. Wound closure was quantified as the percentage of healed area relative to the initial scratch area.

### Quantitative real-time PCR (qRT-PCR)

Total RNA was isolated from treated cells using the NucleoSpin® Gel and PCR Clean-Up Kit (Takara Bio USA, cat.# 740984) and reverse transcribed into cDNA using SuperScript IV VILO Master Mix (Thermo Fisher Scientific, cat.# 11756050). qRT-PCR was performed to assess mRNA expression of MMP9 and EMT-related genes. GAPDH was used as an endogenous control, and relative gene expression was calculated using the 2^−ΔΔCt method^82^. All primers were obtained from Thermo Fisher Scientific.

### Gene silencing

E-cadherin knockdown was performed using small interfering RNAs (siRNAs), including a non-targeting control siRNA (Santa Cruz Biotechnology, sc-37007) and E-cadherin-specific siRNA (sc-35242). siRNAs were transfected into TIGK, HSC3, or HN30 cells using Lipofectamine LTX with Plus Reagent (Invitrogen, cat.# 15338), according to the manufacturer’s instructions. Cells were incubated for 72-96 h post-transfection and subsequently treated with InlA or InlB for the indicated times prior to downstream assays. Knockdown efficiency was confirmed by immunoblotting and immuno-fluorescence staining for E-cadherin.

### Immunoblotting and co-immunoprecipitation (co-IP)

Total protein lysates were prepared using Pierce IP Lysis Buffer (Thermo Fisher Scientific, cat.# 87787). Equal amounts of protein were resolved by SDS-PAGE, transferred to PVDF membranes, blocked with 5% skim milk in PBS for 1 h and incubated overnight at 4°C with primary antibodies against β-catenin, AKT, phospho-AKT, p38, phospho-p38, JNK, phospho-JNK, MMP9, SNAIL, SLUG, TWIST, ZEB1, and β-actin (loading control). After washing, membranes were incubated with appropriate secondary antibodies at RT for 1 h. Signals were detected using enhanced chemiluminescence (ECL) and quantified with a ChemiDoc Imaging System and Image Lab software (Bio-Rad).Co-IP was performed as previously described^24,75^. Briefly, His-tagged recombinant InlA or InlB (20 µg/ml) was incubated with TIGK cell lysates (1 mg total protein) for 3 h, followed by overnight incubation with Ni-NTA agarose beads. Beads were washed five times with PBST, and bound proteins were eluted and analyzed by SDS-PAGE and immunoblotting using anti-His and anti-E-cadherin antibodies. In parallel, His-tagged InlA or InlB protein (20 µg/ml) was incubated with recombinant E-cadherin (5 µg) and processed identically to confirm direct interaction.

### β-catenin phosphorylation and nuclear translocation

TIGK cells were treated with recombinant InlA or InlB (20 µg/mL) for 2 h or overnight. Phosphorylation of β-catenin was assessed by immunoblotting using phospho-β-catenin specific antibodies (Thermo Fisher Scientific, cat.# 702969 and PA5-36748). Nuclear translocation of β-catenin was evaluated by immunofluorescence staining using a β-catenin antibody. In parallel, nuclear proteins were isolated using a nuclear extraction kit (Abcam, cat.# ab113474), and nuclear β-catenin levels were analyzed by immunoblotting. Proliferating cell nuclear antigen (PCNA; Thermo Fisher Scientific, cat.#14-9910-82) was used as a nuclear protein loading control.

### CCK-8 cell viability and drug sensitivity assays

Cell viability was assessed using the CCK-8 assay (Abcam, ab228554). For proliferation assays, 5 × 10³ TIGK, HSC3, or HN30 cells per well were seeded into 96-well plates, pretreated with or without recombinant InlA or InlB (20 µg/mL), and cultured for 24, 48, 72, or 96 h. At the indicated time points, CCK-8 reagent was added and incubated for 2 or 4 h, and absorbance was measured at 460 nm wavelength. For drug sensitivity assays, 2.5 × 10⁴ TIGK, HN30, or HSC3 cells were seeded into 96-well plates and cultured overnight. Cells were pretreated with recombinant InlA, InlB, or TDE0471C (20 µg/ml) for 2 h, followed by treatment with cisplatin (0, 1, 3.3, 10, or 20 µg/mL) or 5-fluorouracil (5-FU; 0, 1, 3.3, 10, 20 µg/mL) for 48 h. CCK-8 reagent (10 µL) was then added and incubated for 2 to 4 h. Cell viability was calculated relative to untreated controls (defined as 100%), and half-maximal inhibitory concentration (IC₅₀) values were determined.

### Analysis of apoptosis by Annexin V staining and immunoblotting

Apoptosis was assessed by immunofluorescent staining using Annexin Vstaining^83^. TIGK, HN30, or HSC3 cells (1.5 × 10⁶) were seeded into 6-well plates and cultured overnight, pretreated with recombinant InlA, or InlB (30 µg/mL) for 2 h, and subsequently exposed to cisplatin (10 µg/mL) or 5-fluorouracil (5-FU; 5 µg/mL) for 48 h. Cells were harvested, washed, and stained with Annexin V in binding buffer for 15 min at room temperature in the dark. Immunofluorescent staining analysis was performed to determine the apoptotic cells. In parallel, cell lysates were prepared from treated samples and subjected to immunoblotting analysis to assess apoptosis-related proteins, including cleaved poly(ADP-ribose) polymerase (PARP; Cell Signaling Technology, cat.# 5625), cleaved caspase-3 (cat.# 9661), cleaved caspase-9 (cat.# 9505), Bax (cat.# 2772), and Bcl-2 (cat.# 4223).

### Tumor metastasis studies in mice

HN30-Luciferase cells were pretreated with or without InlA or InlB (20 µg/mL) for 2 h, washed with PBS, and resuspended in PBS. A total of 5 × 10⁵ cells per mouse were injected via the tail vein into NSG mice (Cancer Mouse Models Core, CMMC). Mice were randomly assigned to three groups (n = 6 per group): control (PBS-pretreated cells), InlA- or InlB-pretreated. Tumor growth was monitored weekly for 5 weeks using an IVIS imaging system to detect luciferase activity. At the study endpoint, mice were euthanized, and lungs were collected for histological analysis. Hematoxylin and eosin (H&E) staining was performed to visualize tumor nodules, and nodule number and area were quantified manually and using ImageJ software, respectively.

### Tissue microarrays and immunohistochemical (IHC) staining

A head and neck cancer tissue microarray containing 24 cases (48 cores) with stage and grade information, along with unmatched adjacent normal tissues, was obtained from TissueArray.Com(Cat# HN481). Formalin-fixed, paraffin-embedded sections were deparaffinized in xylene and rehydrated through graded ethanol. Antigen retrieval was performed in citrate buffer (pH 6.0) at 95 °C for 20 min, followed by cooling to room temperature. Endogenous peroxidase activity was quenched with 3% hydrogen peroxide for 15 min, and nonspecific binding was blocked with 5% normal goat serum for 1 h. Sections were incubated overnight at 4 °C with a primary antibody against InlB (generated in-house; 1:300 dilution). Signal was detected using a biotinylated secondary antibody and streptavidin-HRP, followed by visualization with 3,3′-diaminobenzidine (DAB). Slides were counterstained with Mayer’s hematoxylin, dehydrated, and mounted.

### Model preparation and structural comparison

Models of protein complexes between the Homo sapiens E-cadherin-1 ECI domain (Asp155-Phe262, Uniprot ID: P12830) and *P. gingivalis* W83 internalin-related protein (Uniprot ID: Q7MX64, ipTM = 0.49, pTM = 0.76) or immunoreactive 47 kDa antigen (Uniprot ID: F5HDB6, ipTM = 0.35, pTM = 0.74) were generated using AlphaFold3^84^. Structures were visualized, analyzed and figures were prepared using PyMol^85^.

### Statistical analysis

Statistical analysis was performed using GraphPad Prism 9. Data are presented as mean ± S.D. values. Statistical differences between groups were determined by Student t test. P value <0.05 was considered as statistically significant.

